# Nuclear LC3 Interactome Profiling Identifies Clathrin Heavy Chain as a Mediator of Nuclear LC3 Translocation in Trabecular Meshwork Cells

**DOI:** 10.64898/2026.09.22.753497

**Authors:** Myoung Sup Shim, Chien-Chich Chou, Mi Sun Sung, Aleks Grimsrud, Vaibhav Desikan, Nikolai Skiba, Paloma B. Liton

## Abstract

Although MAP1LC3B/LC3B (LC3) is best known as a cytoplasmic marker of autophagosome biogenesis, a substantial pool of LC3 resides in the nucleus and shuttles dynamically between nuclear and cytoplasmic compartments, yet the composition and regulation of this nuclear interactome remain poorly defined. Using mass spectrometry-based proteomic profiling of nuclear GFP-LC3 immunoprecipitates from primary human trabecular meshwork (TM) cells, a mechanosensitive ocular cell type, we identified a reproducible nuclear LC3 interactome enriched for proteins containing LC3-interacting region (LIR) and expanded LIR (xLIR) motifs and reported nuclear localization. Among these, clathrin heavy chain (CLTC) emerged as a previously unrecognized nuclear LC3 partner that localizes to the nucleus and colocalizes with nuclear LC3 puncta. CLTC depletion reduced basal LC3-II levels, consistent with a role in autophagosome biogenesis, and markedly impaired nuclear LC3 accumulation induced by both nuclear export blockade with leptomycin B and cyclic mechanical stretch, without altering total CLTC abundance, indicating that CLTC actively promotes LC3 nuclear translocation rather than serving as a passive scaffold. Strikingly, mechanical stress-induced nuclear LC3 trafficking, but not its basal component, was selectively impaired in TM cells derived from glaucoma patients, despite comparable CLTC levels, pointing to a defect in coupling this trafficking pathway to mechanotransduction rather than in the core transport machinery itself. Together, these findings establish nuclear LC3 trafficking as an actively regulated, CLTC-dependent process linked to cytoskeletal and vesicular machinery, and implicate its dysregulation in mechanically stressed glaucomatous cells, providing a framework for understanding how autophagy intersects with nuclear homeostasis, mechanotransduction, and glaucoma pathogenesis.

## INTRODUCTION

Autophagy is a highly conserved intracellular degradation pathway that preserves cellular homeostasis through the turnover of damaged organelles, protein aggregates, and other cytoplasmic constituents. In canonical macroautophagy, a crescent-shaped isolation membrane, termed the phagophore, expands to sequester cytoplasmic cargo and closes to form a double-membrane autophagosome. The autophagosome subsequently fuses with lysosomes, where its contents are degraded and recycled (1).

A central regulator of this process is microtubule-associated protein 1 light chain 3 (MAP1LC3B, LC3B), a core component of the autophagy machinery and one of the most widely used markers of autophagosome formation. LC3B is a member of the ATG8 protein family, which in mammals comprises the LC3 subfamily (LC3A, LC3B, LC3C) and the GABARAP subfamily. These ubiquitin-like proteins coordinate key steps in autophagosome biogenesis, including membrane initiation, elongation, cargo recruitment, and maturation (2). LC3 is synthesized as a precursor that is cleaved by ATG4 proteases to generate LC3-I, a cytosolic form. Upon autophagy induction, LC3-I undergoes conjugation to phosphatidylethanolamine through the coordinated action of the ubiquitinase-like enzymes ATG7 and ATG3, forming LC3-II. This lipidated form associates with both the inner and outer membranes of the growing autophagosome and is widely used as a biochemical and imaging marker of autophagosome formation.

Beyond serving as a marker, LC3 functions as a molecular scaffold that recruits cargo receptors and regulatory proteins through LC3-interacting region (LIR) motifs. The canonical LIR is defined by a short aromatic–hydrophobic core sequence, [W/F/Y]-X-X-[L/I/V], in which the aromatic residue and the hydrophobic residue insert into two conserved hydrophobic pockets on LC3. This core motif is typically embedded within flanking acidic residues that enhance binding affinity through electrostatic interactions (3, 4). Functional LC3-binding sites frequently extend beyond this minimal four-residue core. Expanded or functional LIR motifs (xLIRs) generally comprise a six–amino acid region in which the aromatic residue occupies position 3 and the hydrophobic residue position 6, surrounded by acidic or polar residues that increase binding strength and contribute to ATG8 family selectivity. Post-translational modifications, such as phosphorylation adjacent to the motif, can further enhance affinity by stabilizing electrostatic contacts or promoting favorable conformational changes. This extended framework broadens the spectrum of LC3-interacting proteins and supports context-dependent regulation of LC3-mediated processes (5). Many well-characterized autophagy receptors, including p62, NBR1, OPTN, and NDP52, contain LIR and xLIR motifs that enable them to bridge ubiquitinated cargo to LC3 on the expanding autophagosomal membrane, thereby directing selective autophagy pathways such as mitophagy, aggrephagy, and xenophagy. In addition to cargo receptors, numerous signaling and regulatory proteins harbor LIR motifs, positioning LC3 as a central interaction hub that integrates selective cargo recognition with membrane remodeling and broader cellular signaling networks (5, 6).

LC3 has been classically characterized as a cytoplasmic autophagy protein; however, accumulating evidence indicates that a substantial pool of LC3 resides within the nucleus and dynamically shuttles between nuclear and cytosolic compartments in a tightly regulated manner (7–15). Nuclear LC3 has been detected in association with slowly diffusing nuclear complexes, and its localization appears to be modulated by cellular state (12, 15). Despite this growing recognition, the precise function of nuclear LC3 remains incompletely defined.

One proposed role for nuclear LC3 is that of a regulatory reservoir that modulates LC3 availability and overall autophagic capacity. Under nutrient-rich conditions, LC3 accumulates in the nucleus in an acetylated form. Upon starvation, SIRT1-mediated deacetylation promotes LC3 export to the cytoplasm, where it becomes available for lipidation and incorporation into autophagosomes (8, 9). The shuttling factor TP53INP2/DOR further facilitates LC3 relocalization and autophagy initiation (16, 17), supporting the concept that nuclear LC3 contributes to rapid autophagic activation in response to stress. Beyond serving as a mobilizable reservoir, nuclear LC3 has also been implicated in the selective degradation of nuclear components, broadly termed nucleophagy (18, 19). Lamin B1 undergoes LC3-dependent autophagic degradation during oncogene-induced senescence, directly linking autophagy to nuclear lamina remodeling (20, 21). Autophagy likewise contributes to the clearance of micronuclei and damaged chromatin, thereby supporting genome stability and nuclear quality control (22–24). Additional studies have reported LC3 association with nucleolar complexes and interaction with TP53INP2/DOR, a nucleolar protein involved in rRNA transcription, suggesting that LC3 may participate in surveillance and regulation of nucleolar function (16). Consistent with this concept, our laboratory previously reported enrichment of LC3 in the nucleolus in association with the ribophagy receptor NUFIP1 in trabecular meshwork cells subjected to mechanical stress (15).

To define the molecular network associated with nuclear LC3, we performed mass spectrometry–based proteomic profiling of LC3-associated proteins in isolated nuclear fractions. This unbiased approach enabled systematic identification of nuclear LC3-interacting partners, including potential LIR/xLIR-containing proteins involved in chromatin organization, RNA processing, nuclear envelope dynamics, and stress signaling. Using primary human trabecular meshwork (TM) cells as a physiologically relevant model of mechanotransduction, we identified a reproducible nuclear LC3 interactome enriched for LIR/xLIR-containing and nuclear-localized proteins and uncovered clathrin heavy chain (CLTC) as a previously unrecognized nuclear LC3 partner. We further demonstrate that CLTC is required for both basal and stress-induced nuclear accumulation of LC3, providing mechanistic evidence that LC3 nuclear trafficking is actively regulated and linked to cytoskeletal and vesicular machinery rather than resulting from passive diffusion or serving solely as a static reservoir. Importantly, nuclear LC3 trafficking is dysregulated in glaucomatous TM cells, suggesting that disruption of this regulatory pathway may contribute to impaired nuclear homeostasis and disease-associated TM dysfunction. Together, these findings expand our understanding of nuclear LC3 function and provide a framework for determining whether LC3 acts as a selective adaptor for nuclear cargo degradation, a regulator of nuclear architecture, or a component of broader stress-responsive pathways. Defining these mechanisms will help clarify how autophagy intersects with nuclear homeostasis, mechanotransduction, and disease pathogenesis.

## METHODS

### Reagents

Chemicals and other materials were obtained from the following sources: Bafilomycin A1 (B1793), DMSO (D2650), goat serum (G9023), HEPES (H0887), bovine serum albumin (BSA; A7906), Trion X-100 (Tx1568-1), β-mecaptoethanol (M6250) from MilliporeSigma; phosphate-buffered saline (PBS,10X; Corning, 46-013-CM); Dulbecco’s modified eagle medium (DMEM; 11885084), 100 × penicillin–streptomycin (Pen/Strep; 15240062), fetal bovine serum (FBS; 10082147), 100 × non-essential amino acids (11140050), gentamycin (15750060), Lipofectamine® RNAiMAX Reagent (13778075), DAPI (62248), Fluoromount-G™ (00–4958-02), subcellular protein fractionation kit for cultured cells (78840), protein assay kit (Micro BCA; 23235), SuperSignal™ West Femto Maximum Sensitivity Substrate (34096), Pierce™ ECL western blotting substrate (32209) from Thermo Fisher Scientific; 4 × Laemmli sample buffer (161–747), Precision Plus Protein™ All Blue Prestained Protein Standards (1610373), protein assay dye reagent concentrate (5000006), Blotting-Grade Blocker non-fat dry milk (1706404), Tween 20 (1706531) and polyvinylidene fluoride (PVDF) membrane for protein blotting (1620177) from Bio-Rad; Leptomycin B (Santa Cruz Biotechnology, sc-358688); GFP-Trap®_MA (Chromotek, gtma-20).

### Isolation and maintenance of primary human trabecular meshwork (TM) cells

Primary human TM cells were obtained from TM tissue carefully dissected from donor corneal rims that remained after corneal transplantation procedures performed at the Duke University Eye Center. Cell isolation and culture conditions followed previously established methods (15). Human TM cells were verified based on their characteristic morphology and their induction of myocilin expression following dexamethasone exposure, consistent with established guidelines for TM cell isolation, characterization, and culture (25). Cultures were subcultured at a 1:2 split ratio upon reaching confluence, and passages 4 through 5 were used for all experiments. All procedures involving human tissue adhered to the principles outlined in the Declaration of Helsinki.

### Cyclic mechanical stretch (CMS)

CMS was delivered using a computer-regulated, vacuum-driven Flexcell® FX-5000™ Tension System (Flexcell International Corp., Burlington, NC, USA), following previously published procedures (26). In brief, primary human TM cells were seeded onto type IV collagen–coated flexible-bottom membranes assembled in 6-well BioFlex culture plates (Flexcell International Corp., Burlington, NC, USA) and maintained under static conditions in growth medium until they reached confluence. At that point, the medium was refreshed, and CMS was administered at 20% peak strain (1 Hz) for 24 hours. Parallel control cultures were handled identically but were not subjected to mechanical stretch.

### Isolation of Total and Subcellular Protein Fractions

Cells were rinsed with ice cold PBS and then collected. For total protein extracts, cells were incubated on ice for 30 minutes in modified RIPA buffer (Millipore Sigma, R0278) supplemented at 1:100 with Halt protease and phosphatase inhibitor cocktail (Thermo Fisher Scientific, 78842). Cytoplasmic, nuclear, and membrane enriched fractions were isolated using the Subcellular Protein Fractionation Kit for Cultured Cells (Thermo Fisher Scientific, 78840) following the manufacturer’s protocol. Protein concentrations were determined using either the Micro BCA protein assay kit (Thermo Scientific, 23235) or the Bradford assay with Protein Assay Dye Reagent (Bio-Rad, 5000006).

### Co-Immunoprecipitation

Primary human TM cells were infected with Ad-GFP-LC3 (kindly provided by Dr. Wen-Xing Ding, University of Kansas Medical Center) or Ad-GFP as a control at 20 pfu per cell and maintained for three days. Nuclei were isolated from 2 to 4 × 10⁷ cells and processed as described previously (26). Nuclear pellets were lysed in co-immunoprecipitation buffer containing 20 mM Tris pH 7.5, 137 mM NaCl, 1 mM MgCl₂, 1 mM CaCl₂, 1% NP-40 (IBI Scientific, 9016-45-9), and 10% glycerol. The buffer was supplemented with Halt protease and phosphatase inhibitor cocktail at 1:100 dilution, 50 U/mL benzonase (MilliporeSigma, 70664), and an additional 5 mM MgCl₂. Lysates were incubated with gentle rotation at 4°C for two hours and clarified by centrifugation at 16,000 × g. The resulting supernatants were incubated with GFP-Trap®_MA (Chromotek) at 4°C for 16 hours, and co-immunoprecipitation was completed according to the manufacturer’s instructions.

### Mass spectrometry analysis of immunoprecipitants

Immunoprecipitants were solubilized in 2% sodium dodecyl sulfate, 100 mM Tris-HCl (pH 8.0), reduced with 10 mM DTT, alkylated with 25 mM iodoacetamide, and subjected to tryptic hydrolysis using the HILIC beads SP3 protocol as previously described (27). The protein digests were separated by liquid chromatography using the Vanquish Neo UPLC (Thermo Scientific) and analyzed using Q Exactive HF Orbitrap mass spectrometer (Thermo Scientific) as previously described (28). For label-free protein quantification, raw mass spectral data files (.raw) were imported into Progenesis QI for Proteomics 4.2 software (Nonlinear Dynamics) for duplicate runs alignment of each preparation and peak area calculations. Peptides were identified using Mascot version 2.5.1 (Matrix Science) for searching the UniProt 2023 reviewed human database containing 20,237 entrees. Mascot search parameters were: 10 ppm mass tolerance for precursor ions; 0.025 Da for fragment-ion mass tolerance; one missed cleavage by trypsin; fixed modification was carbamidomethylation of cysteine; variable modifications were oxidized methionine and deamination of asparagine and glutamine. Only proteins identified with 2 or more peptides (FDR < 1%), were included in the protein quantification analysis. Values representing protein amounts were calculated based on a sum of ion intensities for all identified constituent non-conflicting peptides. For protein and peptide normalization between samples, the integrated peak area for each identified peptide was corrected using the factors calculated by automatic Progenesis algorithm utilizing the total intensities for all peaks in each run. Protein abundances were averaged for two duplicate runs for each sample and ratio between control and target immunoprecipitation was calculated. Clustering analysis was performed using the Markov Clustering algorithm implemented in the STRING database. The mass spectrometry proteomics data have been deposited to the ProteomeXchange Consortium via the PRIDE partner repository with the dataset identifier PXD083665 and 10.6019/PXD083665

### Western-blot

For traditional immunoblotting, 2.5 to 5 μg of protein per sample were resolved on 7.5–15% SDS-PAGE gels and transferred to polyvinylidene fluoride membranes. Membranes were blocked in 5% nonfat milk prepared in PBS-T containing 0.1% Tween-20 (Bio-Rad, 170-6531) for one hour at room temperature. Primary antibodies were applied overnight at 4°C, followed by washing in PBS-T and incubation with horseradish peroxidase– conjugated donkey anti-mouse, anti-rabbit, or anti-goat IgG secondary antibodies (1:5,000; Jackson ImmunoResearch, 715-035-151, 711-035-152, or 705-035-147) for two hours at room temperature. Immunoreactive bands were visualized using an enhanced chemiluminescence detection system. Images were acquired and quantified with the ChemiDoc™ Touch Imaging System and Image Lab™ software (Bio-Rad). For automated simple western immunoblotting, protein expression was assessed using the ProteinSimple Jess automated capillary-based Western platform (Bio-Techne, San Jose, CA, USA) with the 12 to 230 kDa separation module (SM-FL004) for all targets. Cell lysates were adjusted to 0.5 µg/µL in 0.1× sample buffer and combined with 1× fluorescent molecular weight standards and 40 mM dithiothreitol to yield a final volume of 5 µL per assay, corresponding to 2.5 µg of total protein. Samples were denatured at 95°C for five minutes before loading into Jess assay plates preloaded with blocking buffer, primary antibodies, horseradish peroxidase–linked secondary antibodies (anti-mouse or anti-rabbit IgG), luminol–peroxide substrate, and wash buffer. Following automated separation and immunodetection, chemiluminescent signals were recorded using Compass for SW version 6.3.0 software (Bio-Techne). Data were presented both as electropherograms, illustrating chemiluminescent peak intensity, and as virtual lane images corresponding to capillary-based signal detection.

Primary antibodies and working dilutions were as follows for conventional western blot: anti-LC3B (1:3,000; Cell Signaling Technology, 3868S), anti-SQSTM1 (1:5,000; MilliporeSigma, P0067), anti-H2B (1:2,000; Cell Signaling Technology, 8135S), anti-TUBA4A (1:5,000; MilliporeSigma, T5168), anti-GFP (1:3,000, Santa Cruz Biotechnology, sc-9996), anti-fibrillarin/FBL (1:3,000; Abclonal, A1136), anti-LAMP1 (1:1,000; Abcam, ab24170), anti-LAMP2 (1:1,000; Santa Cruz Biotechnology, sc-8101), anti-CLTC (1:5,000, Proteintech, 26523-1-AP), and anti-ACTB (1:1,000; Santa Cruz Biotechnology, sc-69879).

Primary antibodies and working dilutions were as follows for automated simple western immunoblotting: GFP (1:50, Santa Cruz Biotechnology, sc-9996), SQSTM1(1:100, MilliporeSigma, P0067), Lamin A/C(1:50, Cell signaling Technology,2032S), NPM1(1:500, proteintech,10306-I-AP), HSPA8(1:200, proteintech,10654-I-AP), CLTC (1:2,500, proteintech,66487-I-Ig), CLINT1(1:50, proteintech,10470-I-AP), MYOC1(1:500, Abclonal, A25204), AP2A1 (1:250, proteintech,29887-I-AP), AP2B1 (1:500, proteintech,15690-I-AP), EPB41L2 (1:50, proteintech,15437-I-AP), EPB41(1:400, proteintech,13014-I-AP), ACTB (1:100; Santa Cruz Biotechnology, sc-69879).

### siRNA transfection and viral transductions

siRNA transfections were performed as previously reported with minor modifications (29). Briefly, primary HTM cells were seeded in 24-well plates and grown to approximately 80% confluence in 0.5 mL of growth medium. After 24 h, cells were transfected with 5 pmol of siRNA targeting CLTC (sc-25067) or with a non-targeting control siRNA (siNC, sc-37007). Transfections were performed using Lipofectamine® RNAiMAX reagent following the manufacturer’s protocol. All siRNAs were purchased from Santa Cruz Biotechnology. Transduction of recombinant adenovirus expressing GFP-LC3 (AdGFP-LC3) or GFP (AdGFP) was performed following established protocols (15). A multiplicity of 10 pfu/cell was used.

### Immunocytochemical analyses

Cells were fixed in 4% paraformaldehyde in PBS for 15∼30 min at room temperature. For CLTC immunostaining, cells were additionally post-fixed for 1 min at room temperature with 100% ethanol pre-cooled to −20°C, followed by three washes with PBS. Cells were then incubated in blocking solution containing 1% goat serum, 2% BSA, 0.1% Trion X-100 in PBS for 30min at room temperature. Primary antibodies were diluted in blocking solution and incubated with the cells overnight at 4°C. After several washes with PBS, cells were incubated for 2 h at room temperature with Alexa Fluor 488 or 594 dye-conjugated goat anti-mouse or rabbit IgG antibodies (Invitrogen, A-11032 or A-11037) diluted 1:1000 in blocking solution. Following washing, nuclei were counterstained with DAPI (1 μg/mL). Images were acquired using a Nikon C2si confocal microscope equipped with a 60×/1.4 NA oil-immersion objective. Confocal z-stacks were collected at 0.2 μm intervals. Images were processed and analyzed using Fiji (ImageJ) and/or Imaris software (version 10.2.0; Oxford Instruments). Primary antibodies used included anti-LC3 (1:1,000; MBL, PM036), anti-ATG16L1 (1:1,000, MBL, PM040) and anti-CLTC (1: 500, Proteintech, 66487-1-Ig).

### Three-dimensional reconstruction and spatial association analysis of nuclear GFP-LC3 and CLTC

Confocal z-stack images were analyzed using Imaris software (version 10.2.0; Oxford Instruments). DAPI, CLTC, and GFP-LC3 signals were reconstructed in three dimensions, and CLTC- and GFP-LC3-positive puncta were rendered as spot objects to visualize their spatial distribution within the nucleus. Three-dimensional reconstructions were examined from multiple viewing angles, and rotational animations were generated. For quantitative spatial association analysis, individual nuclei were defined by DAPI staining, and CLTC and GFP-LC3 signals were restricted to the corresponding nuclear masks. Puncta in each channel were independently detected using the Spots module, and their spatial association was determined using the Colocalize Spots function. A center-to-center distance threshold of ≤0.3 μm was used to define spatial association between CLTC and GFP-LC3 spots, based on previously described Imaris-based spot proximity analysis (30). For each cell, the percentage of CLTC spots associated with GFP-LC3 spots and the percentage of GFP-LC3 spots associated with CLTC spots were calculated. Individual cells were used for quantification, and the number of cells analyzed is indicated in the corresponding figure legends.

### Statistical Analysis

Data are presented as mean ± standard deviation (SD). Statistical analyses were performed using GraphPad Prism 11 (GraphPad Software, San Diego, CA, USA). Comparisons between two groups were performed using paired or unpaired two-tailed Student’s *t*-tests, as appropriate. Comparisons among multiple groups were analyzed using two-way analysis of variance (ANOVA) followed by Tukey’s multiple-comparison test. Differences were considered statistically significant at P < 0.05.

## RESULTS

### Identification and Characterization of Proteins Selectively Co-immunoprecipitating with Nuclear GFP-LC3 compared to GFP in TM Cells

Primary TM cells were transduced with 10 pfu/cell of AdGFP-LC3 or AdGFP. At day 3 post-infection, nuclear fractions were isolated and subjected to GFP-Trap immunoprecipitation (Figure 1A). The purity of the nuclear fractions was verified (Figure 1B). Co-immunoprecipitation quality was further validated by confirming the previously reported association of nuclear LC3 with sequestosome 1 (SQSTM1) and fibrillarin (FBL) (Figure 1C) (15). Immunoprecipitants were analyzed by mass spectrometry. Data are available via ProteomeXchange with identifier PXD083665. A total of 151 proteins were identified that exhibited at least 1.5-fold enrichment in GFP-LC3 samples compared with GFP controls (*q* value < 0.05) and were represented by more than one unique peptide (Table S1). Among these, 54 proteins displayed greater than threefold enrichment (Table 1). As expected, MAP1LC3B was the most highly enriched protein. To further characterize these candidates, the identified proteins were cross-referenced with public databases to assess the presence of LC3-interacting regions (LIRs and high-confidence xLIRs motifs, (5)) and reported nuclear localization (31). 92% of the proteins contained LIR motifs, 73% have reported nuclear localization, and 76% both contain LIR motifs and are localized to the nucleus. Most of the top candidate proteins contained at least one predicted high-confidence xLIR motif, further strengthening our findings. Examples include microtubule-associated protein 1B (MAP1B), microtubule-associated protein 1A (MAP1A), clathrin heavy chain 1 (CLTC), microtubule-associated protein 1S (MAP1S), band 4.1-like protein 2 (EPB41L2), sequestosome 1 (SQSTM1/p62), and forkhead box protein K1 (FOXK1). STRING network analysis of the 54 enriched proteins was conducted to identify clusters of physical interactions (Figure 1D). This analysis revealed several significantly enriched functional modules, including ribosome biogenesis, autophagy, translation, cortical cytoskeleton organization, clathrin-mediated vesicle trafficking, and genome imprinting.

**Figure 1.**
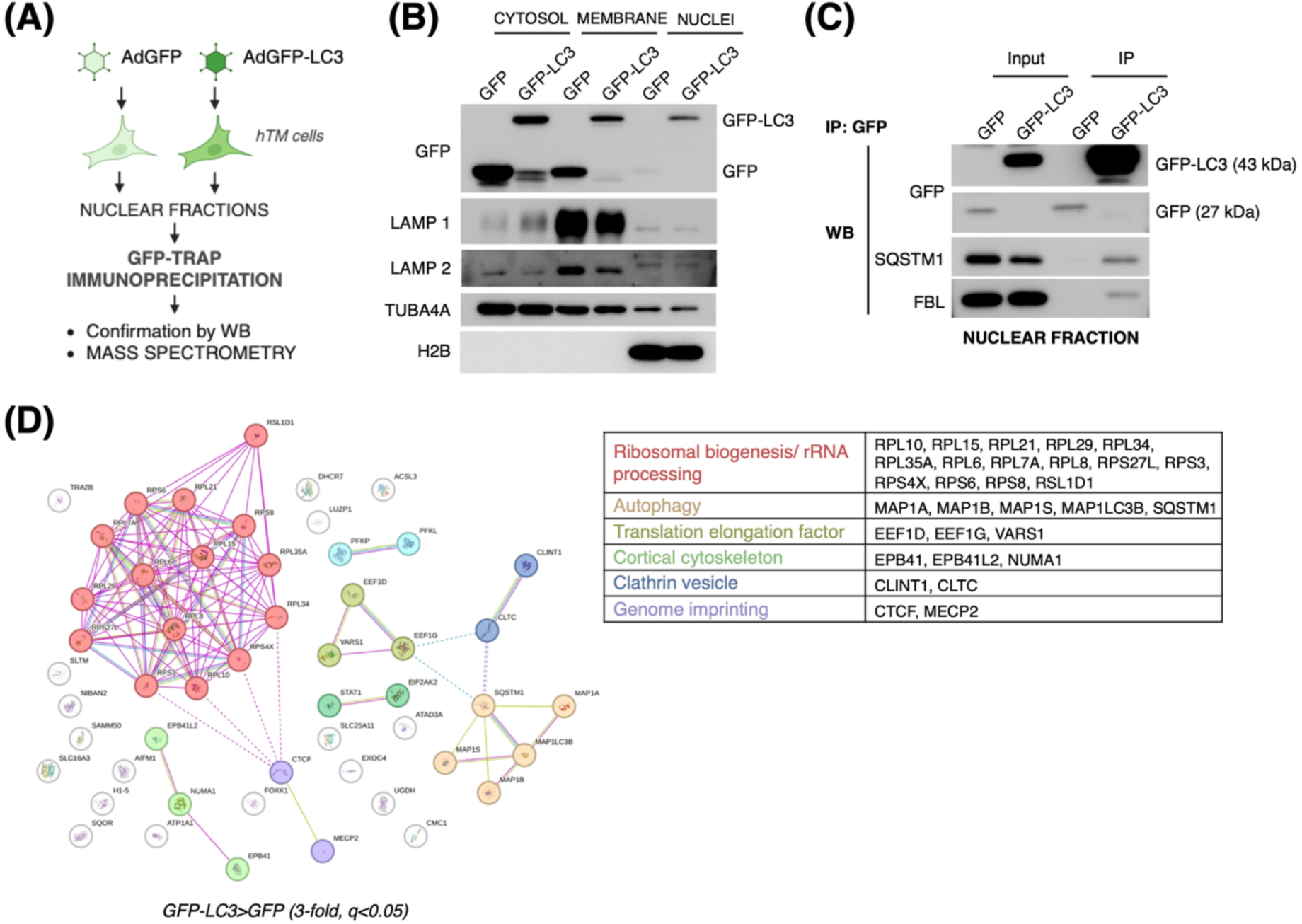
Identification of nuclear LC3-interacting proteins in TM cells. **(A)** Schematic overview of the experimental workflow. **(B)** Subcellular fractionation of AdGFP- and AdGFP-LC3–transduced hTM cells. Cytosolic, membrane, and nuclear fractions were analyzed by WB using antibodies against GFP, LAMP1, LAMP2, TUBA4A, and histone H2B to assess fraction purity. **(C)** Validation of nuclear GFP-LC3 immunoprecipitation. Nuclear extracts were subjected to GFP IP, followed by WB for GFP, SQSTM1, and FBL. Input and IP fractions are shown. **(D)** STRING protein interaction network of the 54 proteins enriched in GFP-LC3 immunoprecipitates. Proteins within each cluster are listed.

**TABLE 1:**
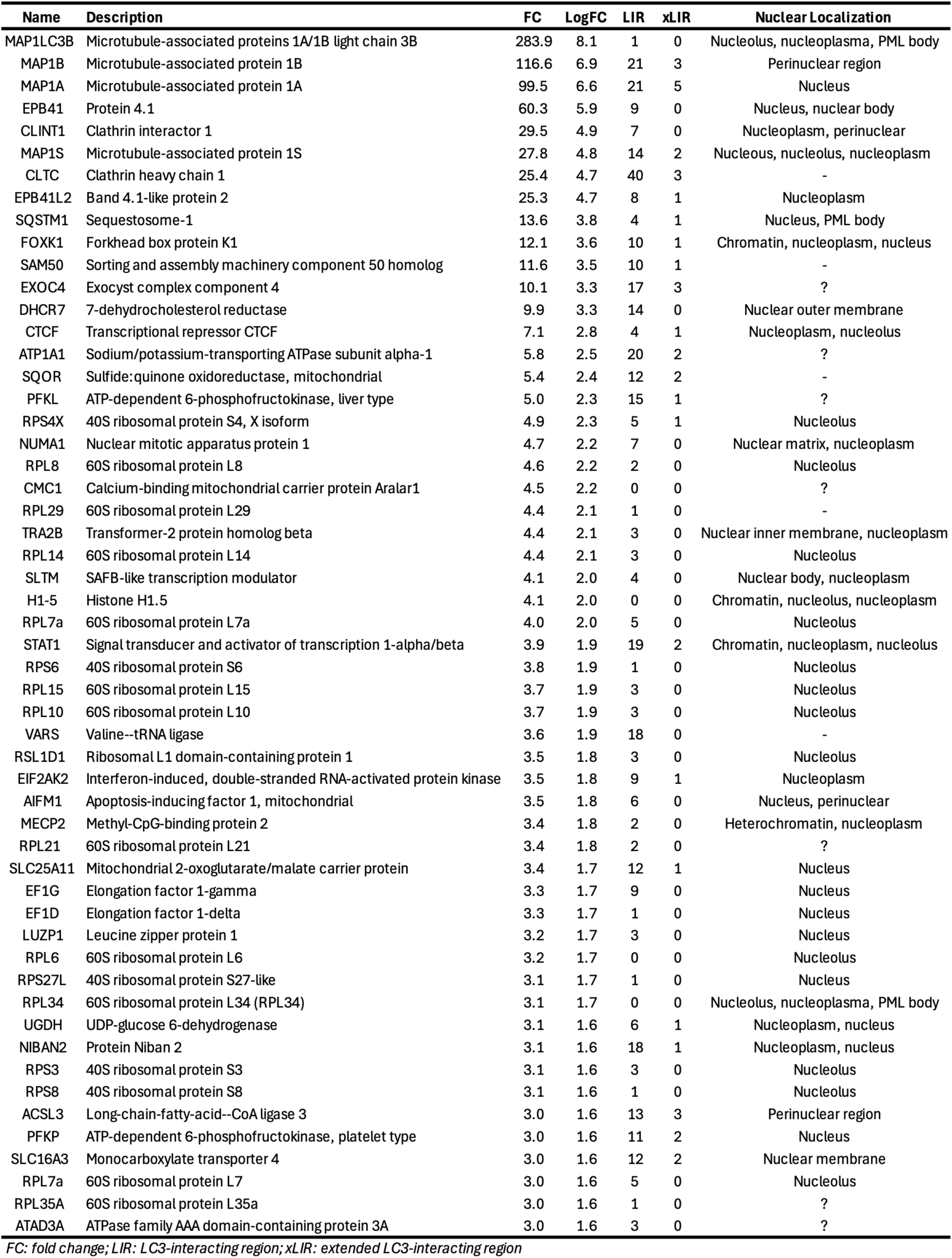
List of proteins co-immunoprecipitating with GFP-LC3 vs GFP (>3-fold, q<0.05, >1 unique petide)

### Identification of Proteins Co-immunoprecipitating with Nuclear LC3 in TM Cells

Our previous work demonstrated increased nuclear LC3 levels in TM cells in response to mechanical stretch. We therefore investigated whether this increase was associated with changes in the nuclear LC3-interacting proteome. For this, we transduced two independent strains of primary TM cells (hTM1, hTM2) with AdGFP-LC3. At 2 days post-transduction, cells were subjected to cyclic mechanical stretch (20 % elongation, 24 h), while control cells were maintained under identical experimental conditions without stretch. Nuclear fractions were isolated, and GFP-Trap immunoprecipitation was performed. Purity of nuclear fractions was confirmed (Figure S1A) and co-immunoprecipitation with SQSTM1 and/or FBL validated the quality of the immunoprecipitation (Figure S1B). Tables S2 and S3 list the proteins co-immunoprecipitated with GFP-LC3 in hTM1 (67 proteins) and hTM2 (172 proteins), respectively, each represented by more than one unique peptide. Of the 67 proteins identified in hTM1 cells, 45 were differentially represented in stretched compared with non-stretched conditions (q<0.05), with 42 showing increased and 3 showing decreased abundance (Table S4). In hTM2 cells, 82 proteins were differentially represented under stretch (q<0.05), including 77 with increased and 5 with decreased abundance (Table S5).

We first compared the lists of proteins represented by more than one unique peptide in non-stretched and stretched hTM1 (Table S2) and hTM2 (Table S3) cells with proteins that significantly bound GFP-LC3, but not GFP (Table S1), to more rigorously identify proteins that consistently interact with nuclear LC3 across all experiments. The comparative analyses are shown in Figure 2A and Table 2. A total of 10 proteins were identified as interacting with nuclear LC3 across all three independent strains, while 33 proteins were detected in two of the three strains. Several additional proteins, marked with asterisks, were also observed in the remaining strain but were represented by only a single unique peptide. Predicted LIR and high-confidence xLIR motifs, as well as reported nuclear localization, are summarized in Table 2. Overall, 88% of the proteins contain LIR motifs, 79% have reported nuclear localization, and 70% both contain LIR motifs and are localized to the nucleus. Strong candidate interactors of nuclear LC3—based on the number or presence of xLIR motifs and nuclear localization—include CKAP4, PLEC, SQSTM1, MAP1A, MAP1B, MYH10, CLTC, CPT1A, RPL23A, RPL7, SAFB, and SLC25A1.

**Figure 2.**
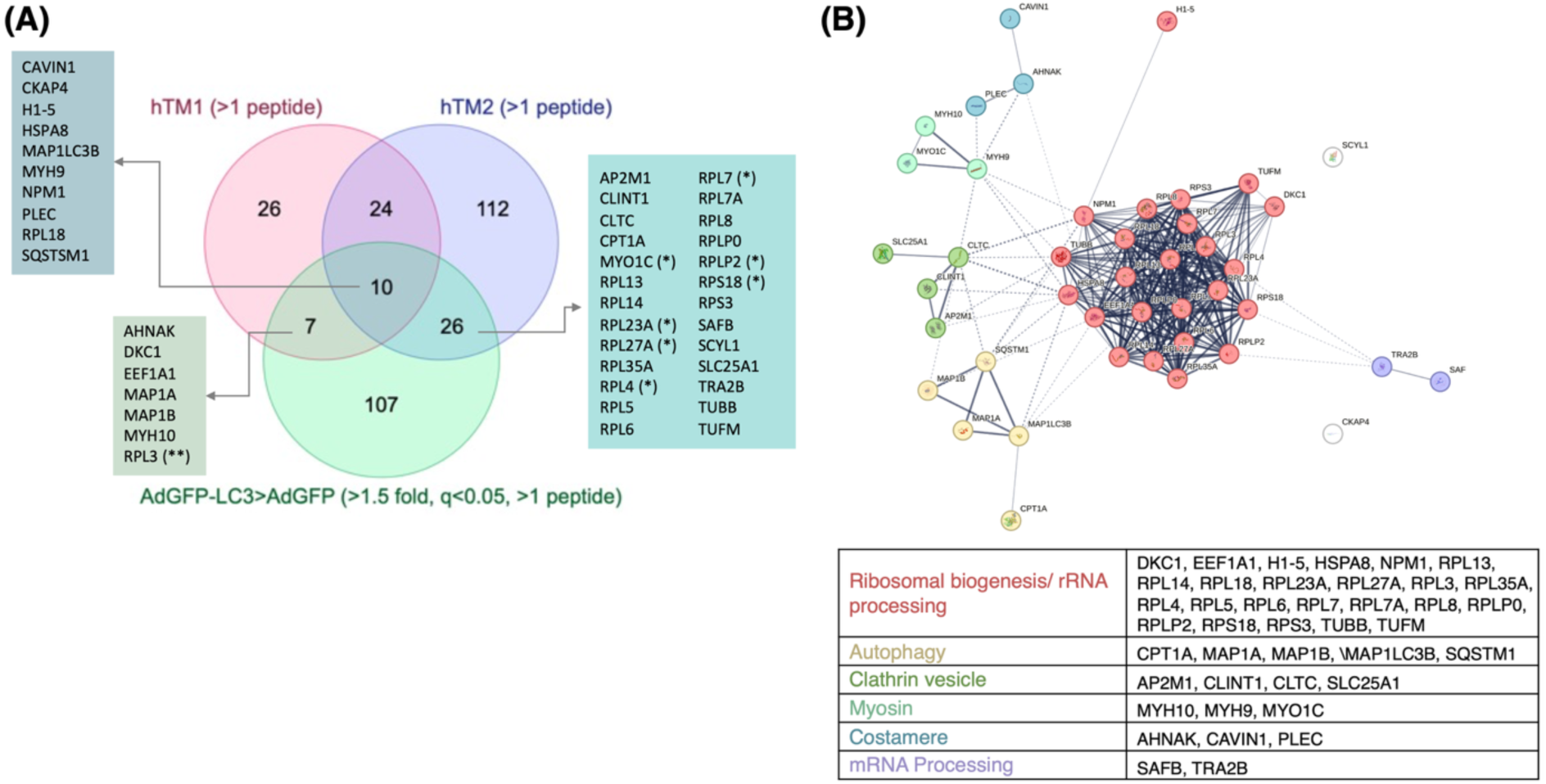
Proteins associated with nuclear GFP-LC3 in non-stretched and stretched hTM cells. **(A)** Venn diagram showing overlap of proteins identified by immunoprecipitation– mass spectrometry as binding partners of nuclear GFP-LC3 across independent experiments in hTM cells. Proteins detected with more than one peptide in hTM1 and hTM2 datasets are indicated (red and blue circles, respectively). The green circle represents proteins enriched in AdGFP-LC3 compared with AdGFP control (>1.5-fold enrichment, q < 0.05, >1 peptide). Numbers indicate the total proteins identified in each dataset and their intersections. Lists at left and right highlight proteins found uniquely or commonly across conditions. **(B)** STRING protein–protein interaction network of proteins consistently detected across experiments. Enriched functional clusters and proteins within each cluster are summarized in the table.

**TABLE 2:**
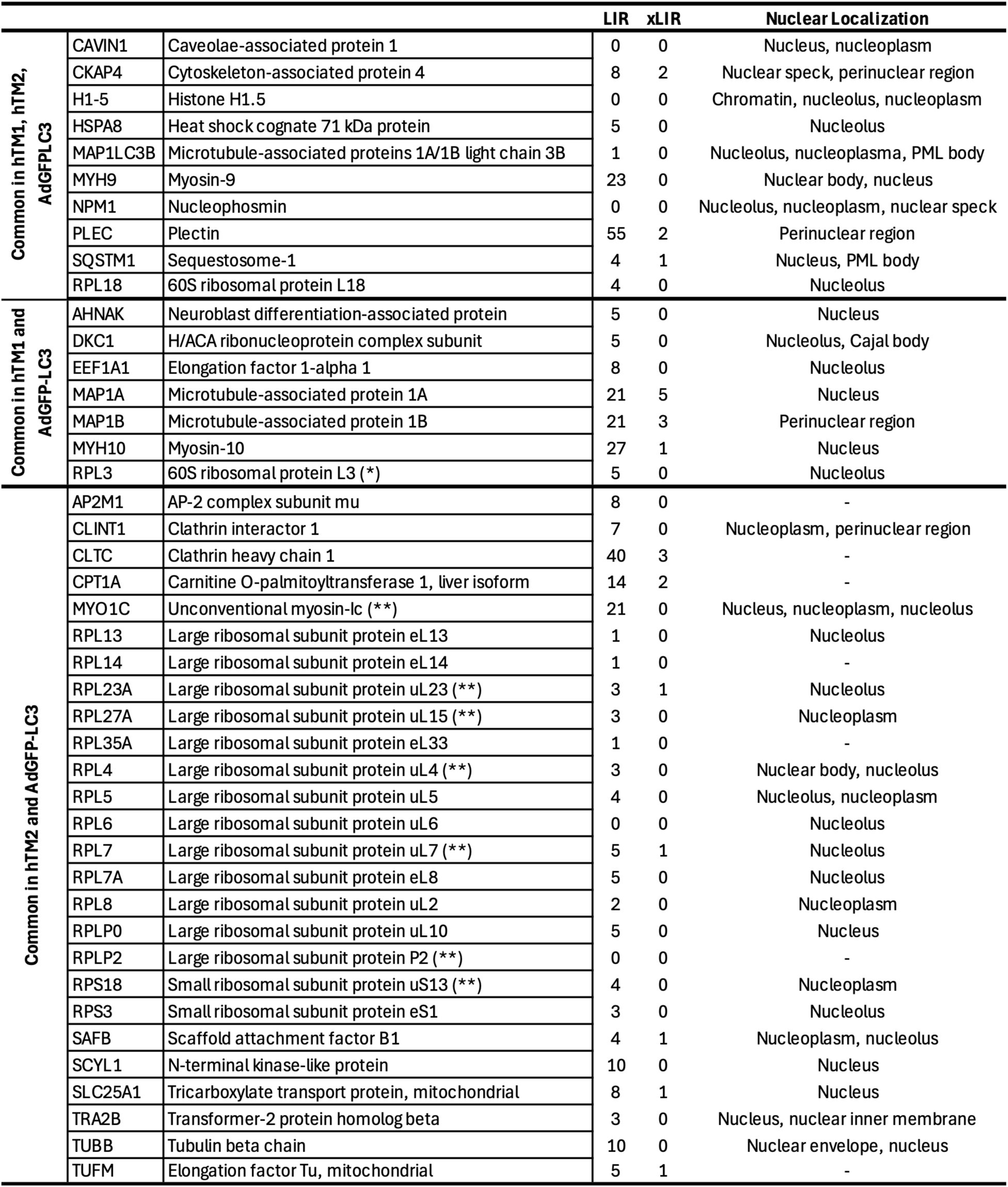
List of proteins consistently co-immunoprecipitating with nuclear LC3 (q<0.05, >1 unique petide)

Full STRING network analysis revealed 6 major enriched pathways (Figure 2B): Ribosome biogenesis, autophagy, clathrin binding, acting binding, costamere, and mRNA processing. An enrichment in nucleolar proteins was also observed, consistent with our prior work (15) and those by others (12) reporting nucleolar localization of LC3.

When comparing the interacting proteome between non-stretched and stretched cells we found no drastic differences, with only 5 proteins showing higher affinity in the stretched cells: CKAP4, HSPA8, NPM1, PLEC and SQSTM1 (Figure S2).

### Comparative analysis of proteins binding GFP-LC3 in cytosolic and nuclear fractions

Several proteins identified as nuclear LC3 interactors, although reported to localize to the nucleus, are constitutively cytosolic proteins that shuttle between the cytosol and the nucleus. We therefore examined whether LC3 also interacts with these proteins in the cytosolic compartment. To address this, we performed GFP-trap co-immunoprecipitation using the soluble cytosolic fraction, excluding membranes and membrane-bound organelles, to identify proteins interacting with LC3 in the cytosol. The complete list of cytosolic LC3-interacting proteins is provided in Table S6. This dataset was compared with the list of proteins immunoprecipitating with LC3 in the nucleus (Table 2). As shown in Figure 3A, of the 43 total proteins identified, 30 interacted exclusively with LC3 in the nucleus, whereas 13 were detected in both nuclear and cytosolic fractions. Interestingly, STRING analysis revealed spatial compartmentalization of functional clusters. Proteins exclusively interacting with nuclear LC3 grouped into ribosomal biogenesis and myosin-associated clusters, whereas proteins detected in both nuclear and cytosolic fractions clustered within the clathrin vesicle trafficking network (Figure 3B).

**Figure 3.**
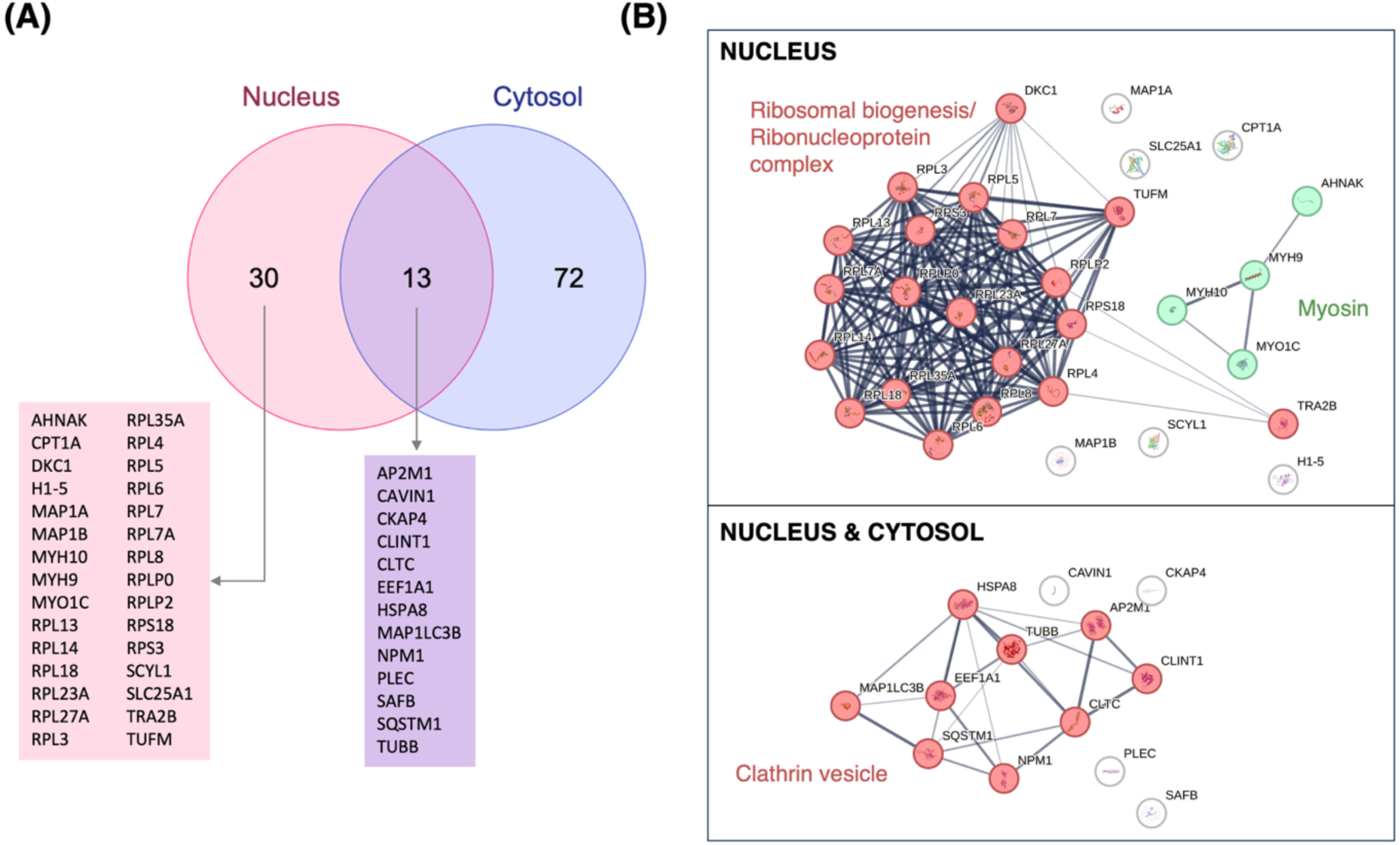
Comparative analysis of proteins binding GFP–LC3 in cytosolic and nuclear fractions. **(A)** Venn diagram showing LC3-interacting proteins identified by mass spectrometry in nuclear and cytosolic fractions. **(B)** Protein–protein interaction networks of LC3-associated proteins. In the nuclear fraction (top), LC3 interactors are enriched in ribosomal biogenesis and ribonucleoprotein complex components (red), with a smaller cluster of myosin-related proteins (green). In the shared nuclear and cytosolic fraction (bottom), LC3-associated proteins form a network enriched in clathrin vesicle trafficking components, including CLTC and adaptor proteins.

We next validated selected protein interactions by Western blot (Figure 4) using newly prepared immunoprecipitants from cytosolic and nuclear fractions. Co-immunoprecipitation with SQSTM1 served as positive control. MYO1C and CLTC were robustly confirmed as LC3-interacting partners in both cytosolic and nuclear fractions, whereas no signal was observed in the GFP control. Additional candidates previously identified by mass spectrometry were detected as LC3-interacting proteins in the cytosol but not in the nucleus, likely reflecting their lower abundance in the nuclear fraction.

**Figure 4.**
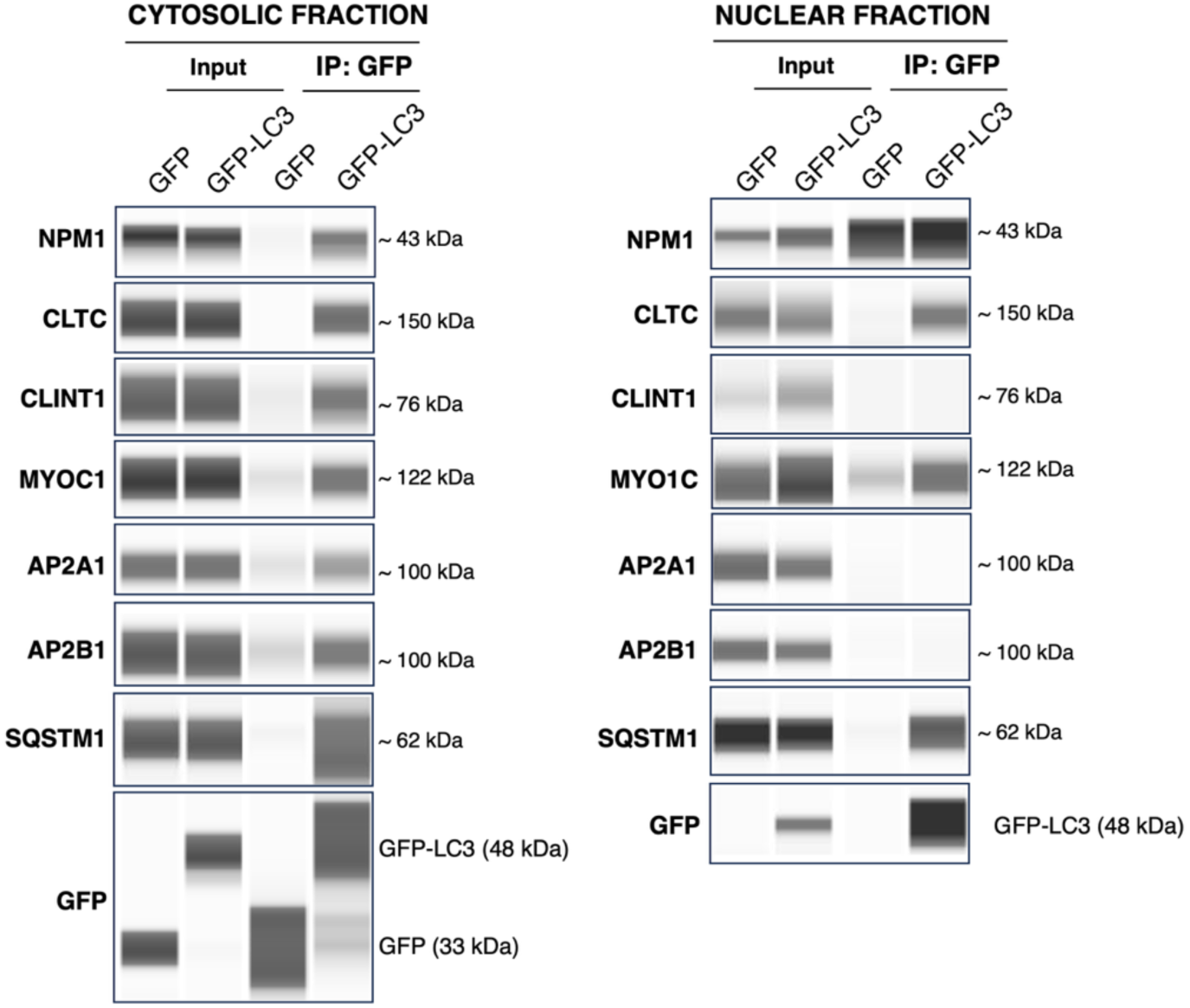
Western-blot validation of LC3-interacting proteins in cytosolic and nuclear fractions. (**A)** Cytosolic and **(B)** nuclear fractions of hTM cells transduced with AdGFP or AdGFP-LC3 were subjected to immunoprecipitation using GFP-trap. Input and immunoprecipitated (IP) fractions were probed by WB for the indicated proteins. SQSTM1 was included as a positive control. GFP and GFP–LC3 are shown as controls for expression and immunoprecipitation efficiency.

### CLTC localizes in the nucleus and interacts with nuclear GFP-LC3

CLTC is a core component of clathrin-coated vesicles that mediate endocytosis and membrane trafficking. Although predominantly cytosolic, monomeric clathrin can translocate to the nucleus, where it performs non-endocytic functions (32). In this context, CLTC has been implicated in chromatin organization, mitotic spindle stabilization (33, 34), and the transactivation of p53 target genes (35). In addition, CLTC has been shown to colocalize with LC3B in the cytosol upon autophagy induction via its xLIR motif (36). However, whether CLTC interacts with nuclear LC3 remains unknown. To address this, we first confirmed the presence of CLTC in the nucleus and assessed its potential association with LC3. TM cells transduced with AdGFP or AdGFP-LC3 were subjected to co-immunolabeling with anti-CLTC antibodies. As shown in Figure 5, CLTC was primarily localized in the cytosol, as expected, but was also detected in the nucleus, where it partially colocalized with nuclear GFP–LC3 (Figure 5B; Supplemental Video).

**Figure 5.**
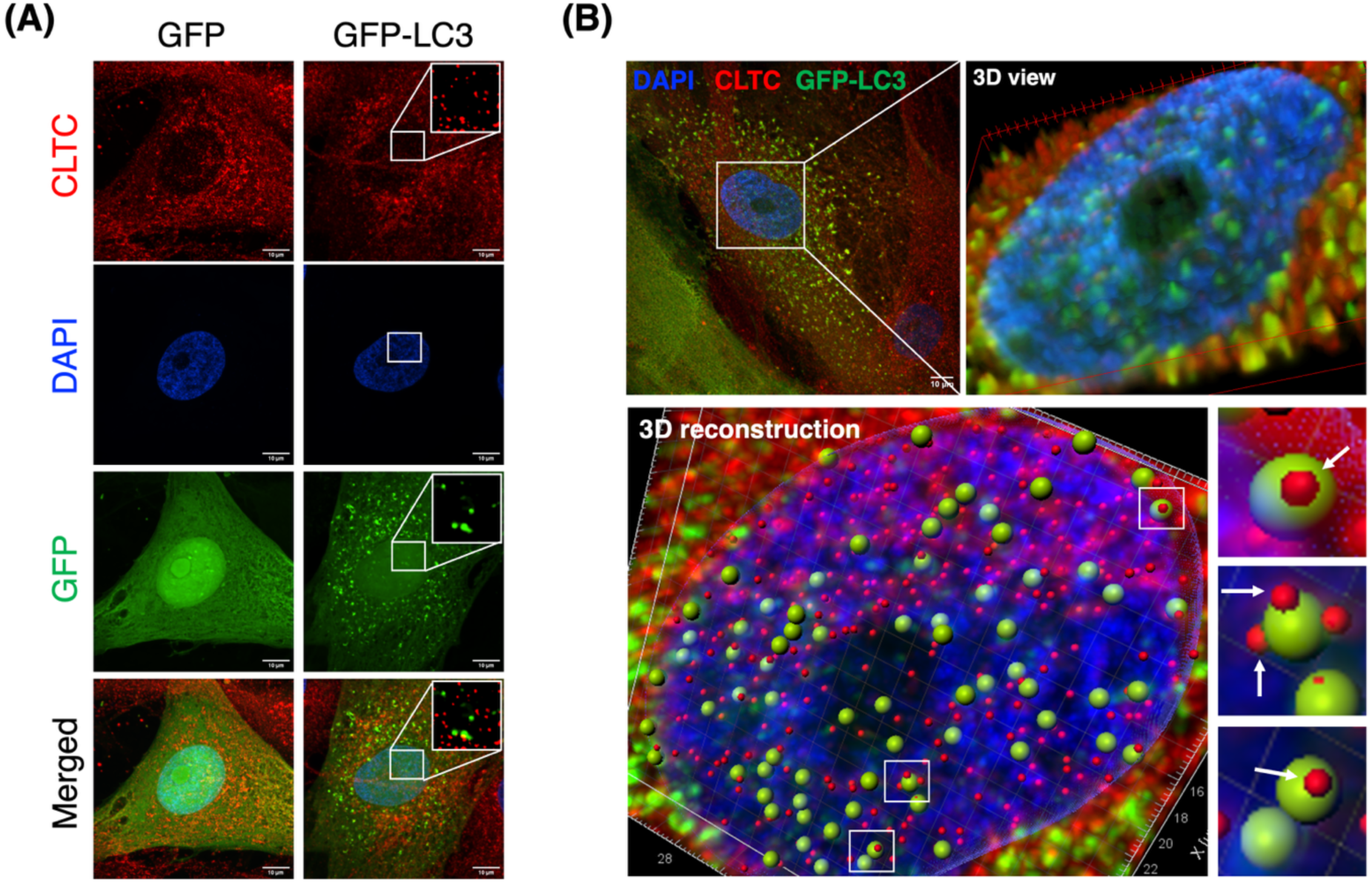
CLTC localizes to the nucleus and associates with GFP-LC3. **(A)** Representative confocal images of cells expressing GFP or GFP-LC3. CLTC (red), nuclei stained with DAPI (blue), and GFP or GFP-LC3 (green) are shown. GFP-LC3 forms puncta that partially co-localize with CLTC (yellow in merged images). Insets highlight overlap. Scale bars, 10 µm. **(B)** High-resolution imaging and 3D analysis of CLTC and GFP-LC3 distribution. Top left: merged confocal image showing DAPI (blue), CLTC (red), and GFP-LC3 (green), with a boxed nuclear region. Top right: 3D view of the nucleus illustrating spatial distribution of signals. Bottom: 3D reconstruction of the nuclear volume showing GFP-LC3 puncta (green) and CLTC signal (red), revealing close spatial proximity and partial overlap. Right panels: magnified views of selected regions demonstrating CLTC-associated GFP-LC3 puncta. Scale bars, 10 µm.

### CLTC is involved in autophagosome formation in TM cells

Previous studies have shown that CLTC is required for both early autophagosome formation and later stages of the process, including regulation of autophagic lysosomal reformation (37–40). Based on these findings, we investigated whether CLTC contributes to autophagy in our cells. For this, we examined LC3-II levels under basal conditions and in the presence of bafilomycin A1 (BafA1, 2h) to assess autophagic flux. As shown in Figure 6A, CLTC silencing led to reduced LC3-II levels, as determined by Western blot, both in the absence of and following BafA1 treatment, suggesting a role for CLTC in autophagosome biogenesis. Immunofluorescence analysis further revealed punctate cytoplasmic distributions of ATG16L1 and CLTC, with partial colocalization (Figure 6B, arrows). CLTC also colocalized with endogenous LC3-positive puncta, preferentially in the perinuclear region. These observations are consistent with prior findings demonstrating that clathrin heavy chain interacts with ATG16L1 facilitating autophagosome formation (37, 39).

**Figure 6.**
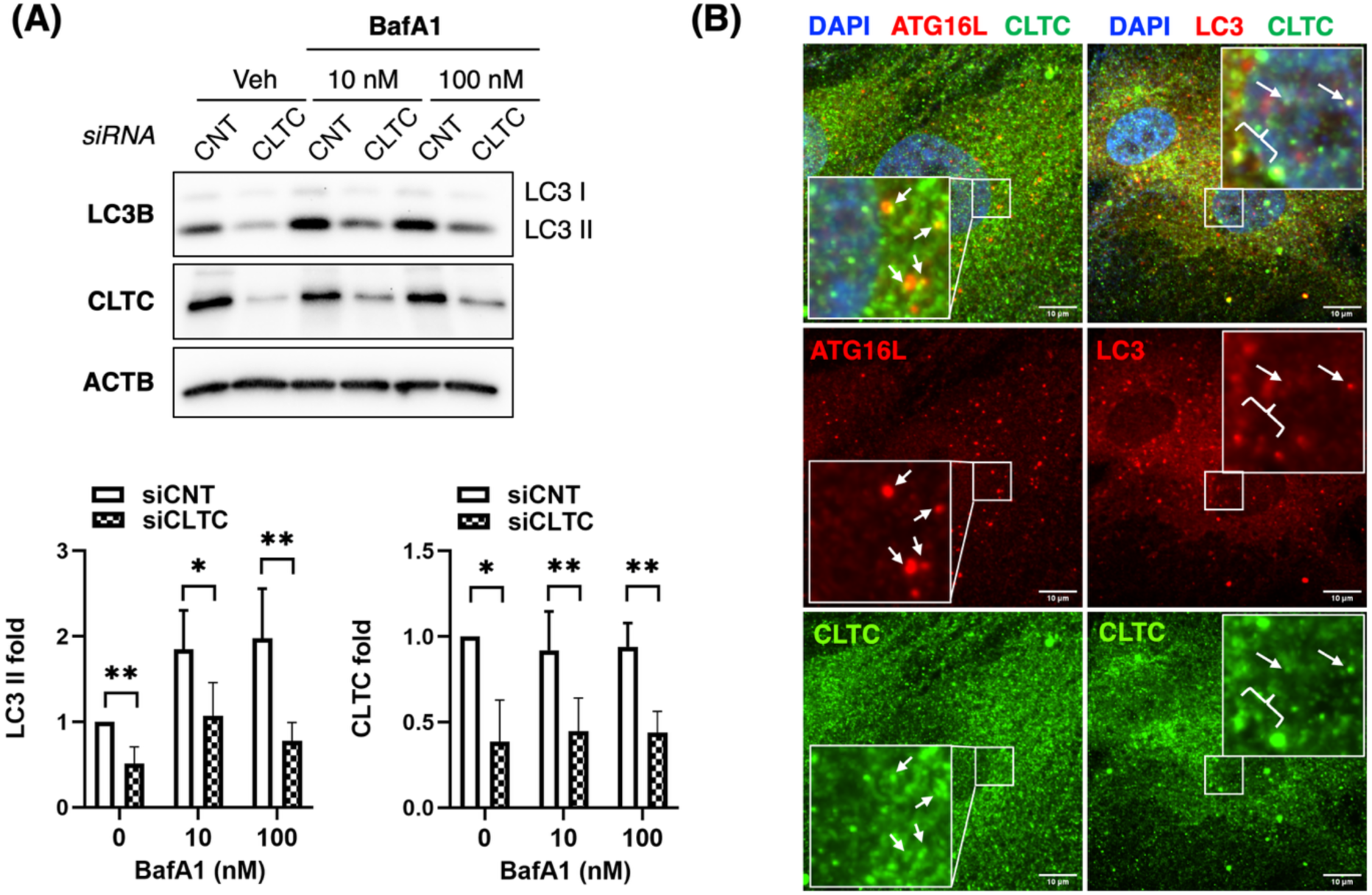
CLTC is required for basal autophagy in human TM cells. **(A)** Human TM cells were transfected with control siRNA (siCNT) or CLTC-targeting siRNA (siCLTC) for 3 days and then treated with vehicle or bafilomycin A1 (BafA1; 10 or 100 nM) for 2 h. Representative immunoblots show LC3B, CLTC, and ACTB protein levels. ACTB was used as a loading control. Densitometric quantification of LC3-II and CLTC protein levels is shown as fold change relative to vehicle-treated siCNT cells. CLTC knockdown reduced LC3-II levels under basal conditions and following BafA1 treatment. Data represent six measurements from three independent hTM cell strains, with two technical replicates per strain. Statistical significance between siCNT and siCLTC at each BafA1 concentration was determined using a paired *t*-test. \**P* < 0.05; \*\**P* < 0.01. **(B)** Representative immunofluorescence images of hTM cells stained for ATG16L1 or LC3 (red), CLTC (green), and nuclei with DAPI (blue). Merged images show partial colocalization of CLTC with ATG16L1-positive and LC3-positive puncta, indicated by arrows and enlarged in the insets. Scale bars, 10 µm.

### CLTC knockdown decreases leptomycin B-induced nuclear accumulation of LC3

Because proteins detected in both the nuclear and cytosolic fractions clustered within the clathrin vesicle trafficking network, we next asked whether CLTC contributes to the nuclear transport of LC3. We assessed GFP-LC3 localization following CLTC knockdown (siCLTC), with or without leptomycin B (LepB, 20 nM for 24 h), a treatment that blocks nuclear export and, as we previously reported, drives LC3 accumulation within PML bodies (15). As anticipated, LepB treatment in control cells (siCNT) led to a pronounced accumulation of GFP-LC3–positive puncta within the nucleus, not observed in cells expressing GFP alone, indicative of LC3 enrichment in PML-associated nuclear structures. Strikingly, depletion of CLTC significantly impaired LepB-induced nuclear accumulation of GFP-LC3 (Figure 7A). Quantitative analysis demonstrated that the percentage of cells containing nuclear GFP-LC3 vesicles was significantly decreased in CLTC-depleted cells compared to controls following LepB treatment (approximately 70–80% in siCNT versus ∼20–30% in siCLTC; ****, p < 0.0001) (Figure 7B). We confirmed these findings biochemically by subcellular fractionation followed by Western blotting. Cytosolic, nuclear, and membrane/organelle fractions were validated using TUBA4A, H2B, and LAMP1, respectively, and efficient CLTC depletion was observed across the fractions (Figure 7C). In siCNT cells, LepB treatment markedly increased nuclear LC3-II levels to approximately 3.3-fold relative to vehicle-treated controls (**, p < 0.001). In contrast, CLTC-depleted cells exhibited only a modest, nonsignificant increase in nuclear LC3-II following LepB treatment. Together, these findings indicate that CLTC is required for efficient LepB-induced accumulation of LC3 within the nucleus.

**Figure 7.**
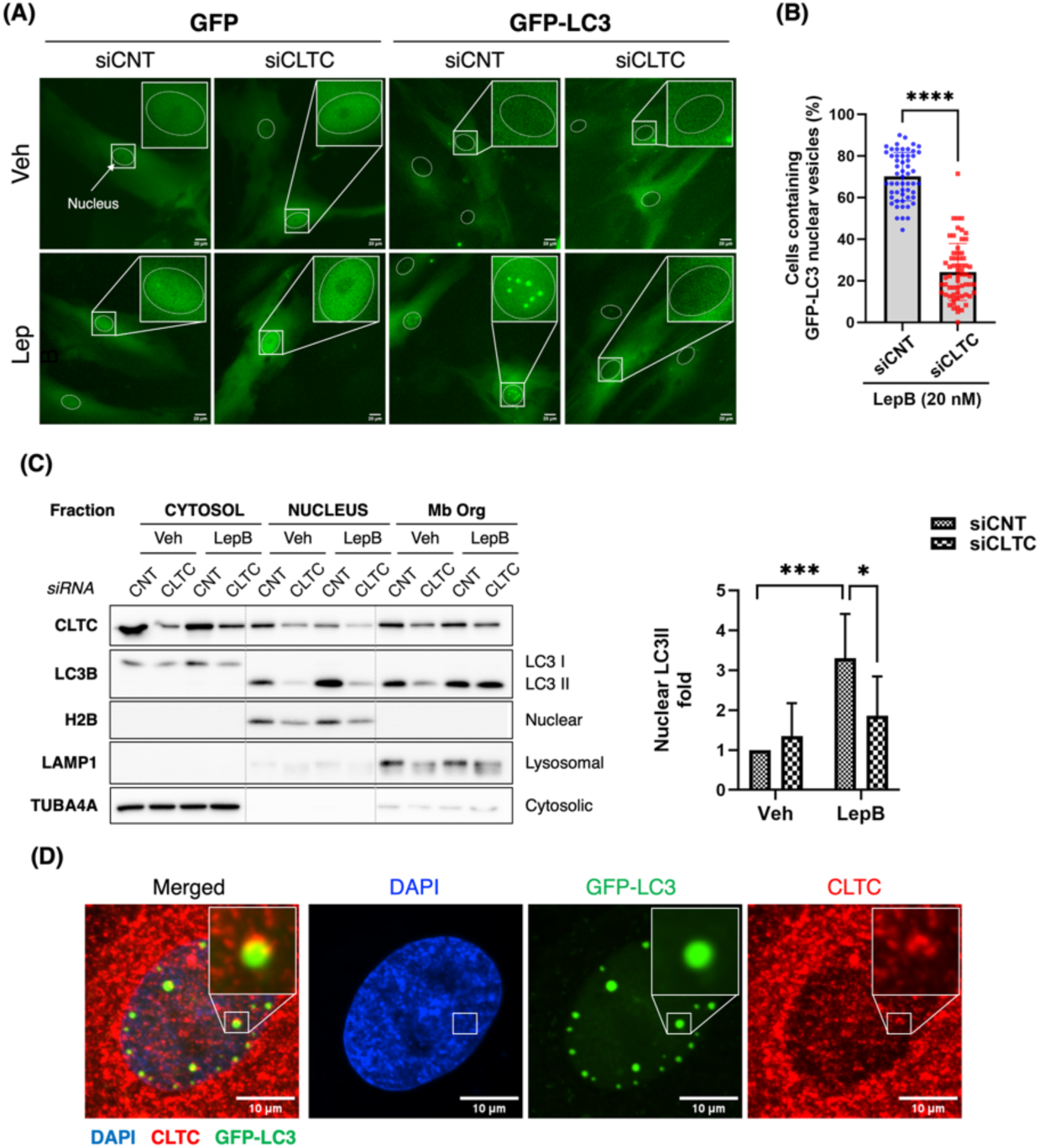
CLTC knockdown reduces leptomycin B-induced nuclear GFP-LC3 vesicles in human TM cells. **(A)** Representative fluorescence images of hTM cells expressing GFP or GFP-LC3 following transfection with control siRNA (siCNT) or CLTC-targeting siRNA (siCLTC) for 3 days. Cells were subsequently treated with vehicle or leptomycin B (LepB; 20 nM) for 24 h. Dashed outlines indicate nuclei, and boxed regions are shown at higher magnification. Scale bars, 20 μm. **(B)** Quantification of the percentage of LepB-treated cells containing nuclear GFP-LC3 vesicles. CLTC knockdown significantly reduced the proportion of cells with nuclear GFP-LC3 vesicles compared with siCNT. Data were collected from 55 siCNT fields and 63 siCLTC fields. Bars represent mean with error bars. \*\*\**P* < 0.0001. **(C)** Subcellular fractionation and immunoblot analysis of hTM cells transfected with siCNT or siCLTC for 3 days and subsequently treated with vehicle or leptomycin B (LepB; 20 nM) for 24 h. Cytosolic, nuclear, and membrane/organelle (Mb Org) fractions were analyzed for CLTC and LC3B. TUBA4A, H2B, and LAMP1 were used as cytosolic, nuclear, and lysosomal fraction markers, respectively. Quantification of nuclear LC3-II showed that LepB increased nuclear LC3-II accumulation in siCNT cells, whereas this response was attenuated by CLTC knockdown. Bars represent mean with SD. ns, not significant; *P < 0.05, *P* < 0.01, and *P < 0.001.

To investigate whether CLTC associates with nuclear LC3-positive structures, we examined the spatial relationship between endogenous CLTC and GFP-LC3 in cells treated with LepB. Immunofluorescence staining demonstrated colocalization between CLTC and GFP-LC3– positive nuclear puncta (Figure 7D). Together, these findings suggest that CLTC contributes to the nuclear accumulation of LC3 following inhibition of nuclear export, potentially by facilitating its translocation to the nucleus and/or its retention within nuclear compartments such as PML bodies.

### CLTC promotes nuclear accumulation of LC3 during mechanical stress

Prior reports from our laboratory demonstrated increased nuclear LC3 in mechanically stretch cells. In view of our findings, we next explored a potential role of CLTC in the stretch-induced nuclear translocation of LC3. Immunofluorescence analysis showed enriched co-localization of CLCT and LC3 in the perinuclear region following mechanical stretch suggesting that mechanical stress promotes the association of the two proteins as LC3 traffics toward and into the nucleus (Figure 8A). Quantification also confirmed that CMS significantly increased the proportion of nuclear LC3 puncta colocalized with CLTC, from 14.91 ± 4.89% under NS conditions to 20.73 ± 5.22% following CMS (n = 30, paired t-test, p < 0.001; Figure 8B). More interestingly, CLTC silencing significantly reduced nuclear LC3-II under basal and stretched conditions (Figure 8C, p < 0.05 and p < 0.01, respectively). Together, these findings indicate that CMS enhances the association of CLTC with nuclear LC3-positive structures and that CLTC is required for both basal and stress-induced accumulation of LC3-II in the nuclear compartment of hTM cells.

**Figure 8.**
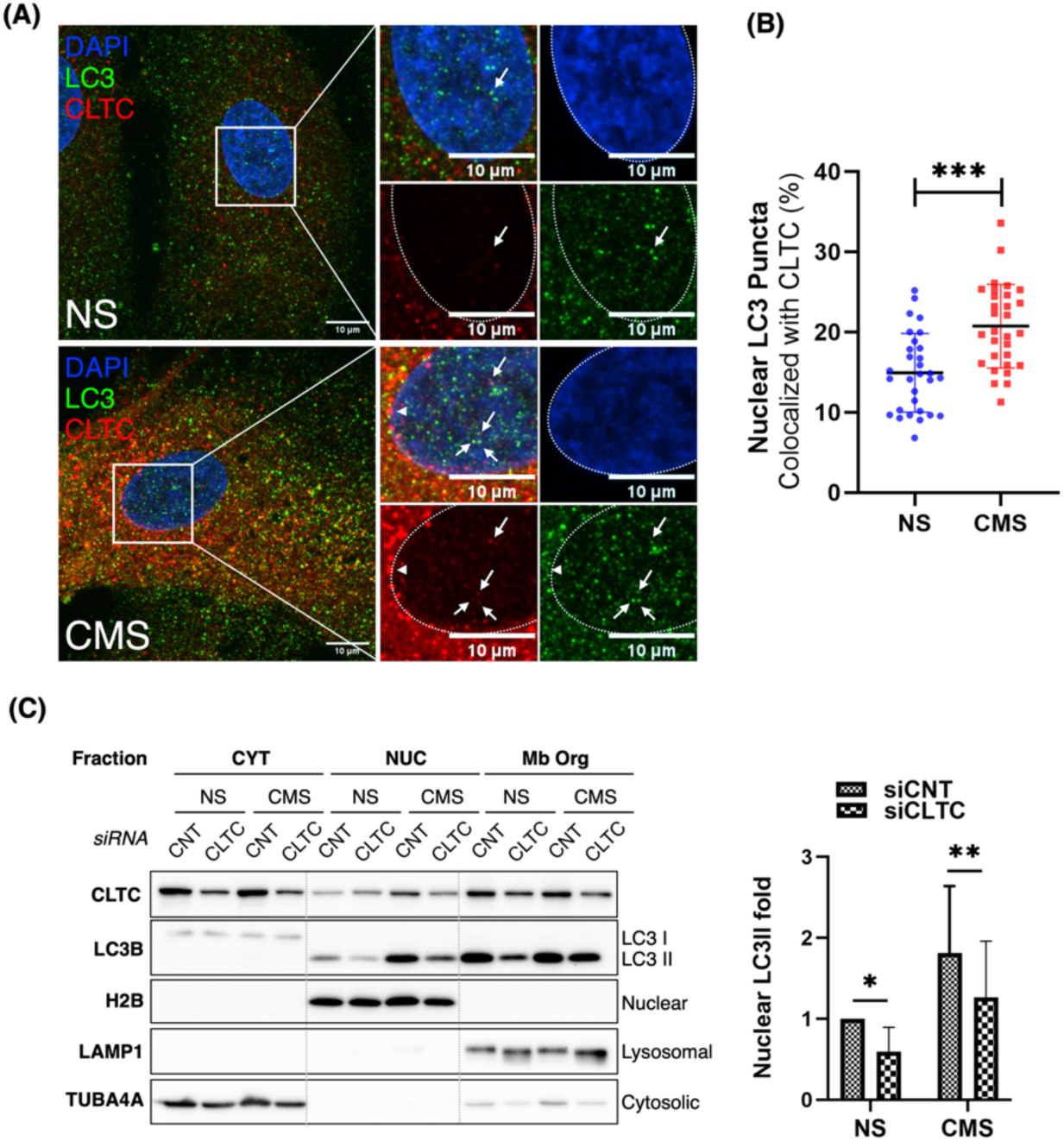
CLTC promotes the nuclear accumulation of LC3 during mechanical stress. **(A)** Representative confocal images of cells maintained under non-stressed conditions (NS) or exposed to cyclic mechanical stress (CMS) and stained for DAPI (blue), LC3 (green), and CLTC (red). Boxed regions are shown at higher magnification. Arrow heads indicate CLTC and LC3 co-localization in the perinuclear region. Arrows indicate LC3-positive puncta within DAPI-defined nuclear regions that are associated with CLTC-positive structures. Dashed lines outline the nuclei. Scale bars, 10 µm. **(B)** Quantification of nuclear LC3 puncta associated with CLTC-positive structures in NS- and CMS-treated cells. Each point represents an individual cell; horizontal lines and error bars indicate mean ± SD. CMS increased the percentage of CLTC-associated nuclear LC3 puncta from 14.91 ± 4.89% to 20.73 ± 5.22%. \*\**P* < 0.001, paired two-tailed Student’s *t*-test. **(C)** Cytosolic (CYT), nuclear (NUC), and membrane/organelle (Mb Org) fractions from control siRNA-treated (siCNT) or CLTC-depleted (siCLTC) cells maintained under NS or CMS conditions were analyzed by immunoblotting for CLTC and LC3B. TUBA4A, histone H2B, and LAMP1 were used as markers for the cytosolic, nuclear, and membrane/organelle fractions, respectively. The graph shows nuclear LC3-II abundance normalized to the corresponding control condition. CLTC depletion reduced nuclear LC3-II accumulation under both NS and CMS conditions. Data are mean ± SD, *P* < 0.05; \**P* < 0.01, paired two-tailed Student’s *t*-test, n=6.

### Mechanical stress selectively impairs LepB-induced nuclear LC3 accumulation in glaucomatous TM cells

Because inappropriate TM adaptation to mechanical stress is thought to contribute to glaucoma pathogenesis, we next asked whether the nuclear LC3 trafficking pathway described above is altered in glaucomatous TM cells. Normal (nTM) and glaucomatous (gTM) TM cell strains (n = 3 each) were maintained under non-stressed (NS) or cyclic mechanical stress (CMS). To block LC3-II nuclear export and thus calculate total LC3-II imported with CMS, we treated the cultures with either vehicle or LepB (20 nM, 24 h). Total nuclear LC3-II levels were quantified by Western blot (Figure 9A). Under NS conditions, total LC3-II levels (LepB-treated cells) did not differ significantly between nTM and gTM strains. As expected, LepB increased nuclear LC3-II in all four groups relative to vehicle (Figure 9B). Interestingly, the total amount of LC3-II translocated in response to CMS in gTM cells was significantly lower in gTM cells compared to nTM cells (Figure 9C). Total CLTC levels did not differ between nTM and gTM strains or between NS and CMS conditions in any subcellular fraction (Figure S3), arguing against a simple expression-level explanation for this difference. Together, these results indicate a mechanical stress–specific, rather than baseline, defect in nuclear LC3 trafficking in glaucomatous TM cells.

**Figure 9.**
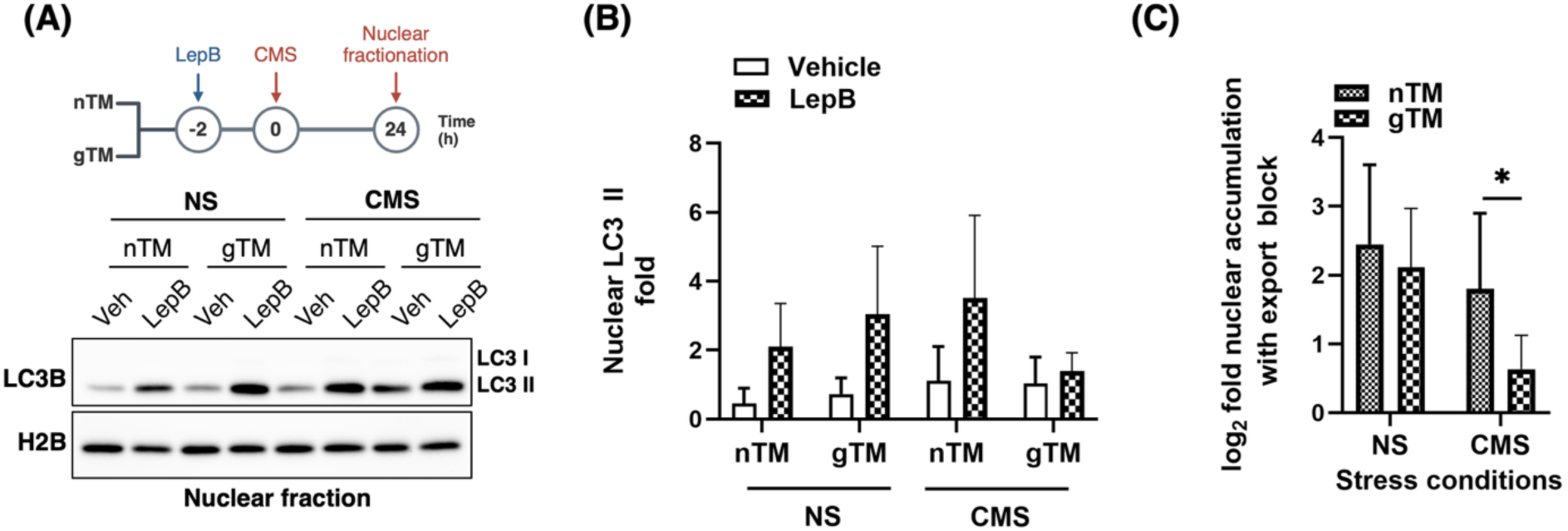
Glaucomatous TM cells exhibit reduced nuclear LC3-II responsiveness to LepB under CMS. **(A)** Experimental schematic and representative immunoblot. Normal (nTM) and glaucomatous (gTM) TM cells were treated with vehicle or LepB (20 nM), added 2 h before the onset of cyclic mechanical stress (CMS) or non-stressed (NS) culture; nuclear fractions were isolated 24 h later and analyzed by WB for LC3B and H2B (nuclear loading control). **(B)** Relative quantification of nuclear LC3-II for each condition. Bars represent mean + SD. **(C)** Nuclear LC3 accumulation following inhibition of nuclear export in nTM and gTM cells under non-stressed (NS) or chronic mechanical stress (CMS) conditions. Nuclear LC3 levels were normalized to H2B, and nuclear accumulation was calculated for each matched experiment as the ratio of normalized LC3 in leptomycin B (LepB)-treated cells to vehicle-treated cells. Data are expressed as log2(LepB/Veh), such that 0 represents no change, 1 represents a 2-fold increase, and 2 represents a 4-fold increase in nuclear LC3 following inhibition of nuclear export. Bars represent mean ± SD from three independent experiments. Statistical analyses were performed on log2-transformed LepB/Veh ratios. The prespecified CMS × genotype interaction was assessed using a paired difference-in-differences contrast (P = 0.048). Pairwise comparisons were performed using paired, two-tailed t-tests with Bonferroni correction for multiple comparisons.

## DISCUSSION

The present study provides a detailed characterization of the nuclear LC3 interactome, supporting a model in which LC3 functions within nuclear protein networks rather than serving as a passive reservoir. Proteomic profiling of nuclear GFP-LC3 immunoprecipitates identified a broad set of associated proteins enriched for predicted LIR/xLIR motifs and nuclear localization signals, indicating selective and functionally relevant interactions in the nuclear compartment. This enrichment was reproducible across GFP-LC3 versus GFP comparisons and multiple independent TM strains under both basal and stretched conditions. Cross-dataset analysis identified a subset of proteins consistently detected in all experiments, alongside a larger group shared between datasets, likely reflecting a combination of biological variability, protein abundance, and technical limitations of mass spectrometry. Proteins repeatedly identified across conditions therefore represent a high-confidence group of nuclear LC3-associated candidates.

A defining feature of this interactome is the marked enrichment of LIR-containing proteins, with the majority of candidates harboring predicted LIR motifs and a substantial fraction also containing high-confidence xLIR sequences. Given the role of LIR motifs in mediating direct interactions with ATG8 family proteins, these findings indicate that many of the identified associations likely reflect direct LC3 binding events (3, 5, 6, 41).The overlap between LIR/xLIR motif prediction and nuclear localization further supports the specificity of these interactions and suggests that canonical ATG8-binding mechanisms extend to the nuclear environment, in agreement with previous studies (10).

Pathway enrichment analysis revealed a strong association of nuclear LC3-interacting proteins with ribosome biogenesis, rRNA processing, chromatin organization, and cytoskeletal-related structures such as clathrin vesicles, myosin complexes, and costameres. The prominence of ribosomal and nucleolar components aligns with prior observations of nucleolar LC3 localization (12, 15), and points toward a functional connection between LC3 and nucleolar homeostasis. These associations are compatible with roles in ribophagy and nucleophagy, processes increasingly recognized as part of nuclear quality control pathways. Supporting this framework, recent studies have demonstrated that autophagy-related mechanisms contribute to the degradation of nuclear components, maintenance of genome integrity, and repair of nuclear damage (18, 19, 22, 42), as well as the regulation of rRNA synthesis (43). In addition, previously reported interactions between LC3 and ribonucleoproteins, including HNRNPK and SAFB, are notable in this context, with SAFB also detected in our dataset, further linking LC3 to ribonucleoprotein complexes and chromatin-associated functions (44).

Comparison of nuclear and cytosolic LC3 interactomes revealed clear compartment-specific interaction patterns, further supported by STRING analysis demonstrating spatial segregation of functional clusters. The limited overlap between nuclear and cytosolic datasets, despite inclusion of soluble cytosolic proteins, suggests that LC3 engages with distinct protein populations depending on its subcellular localization. Intriguingly, the majority of proteins associated exclusively with nuclear LC3 were enriched in ribosome biogenesis and myosin-related processes, whereas shared interactors were predominantly linked to clathrin-mediated vesicle.

Among the identified interactors, CLTC (clathrin heavy chain) emerged as a compelling candidate linking nuclear LC3 function with vesicle trafficking machinery. While clathrin is classically associated with endocytosis and cytoplasmic membrane trafficking, it has also been increasingly implicated in autophagy (37–40). In addition, published evidence supports a functional role for clathrin within the nucleus (32), where it contributes to processes such as chromatin organization and mitotic spindle stabilization (33–35). In this context, our findings extend the known functional repertoire of CLTC by showing that it localizes to the nucleus, interacts with nuclear LC3, and colocalizes with LC3-positive nuclear puncta. Functional analyses further demonstrated that CLTC depletion reduces nuclear LC3-II levels and significantly impairs the accumulation of nuclear LC3 following inhibition of nuclear export and in response to mechanical forces, pointing to a role in coordinating autophagosome biogenesis with nuclear LC3 dynamics.

Our data support a role for CLTC in autophagosome biogenesis in TM cells. CLTC silencing reduced LC3-II levels under both basal conditions and BafA1 treatment, consistent with impaired autophagosome formation rather than a defect confined to downstream degradation. In line with this, CLTC partially colocalized with ATG16L1-positive puncta and with endogenous LC3-positive structures, predominantly in the perinuclear region, placing clathrin at the site where these precursor structures assemble. This functional and spatial link is consistent with two previously described clathrin-dependent routes of autophagosome biogenesis: clathrin heavy chain binds the N-terminal region of ATG16L1 and gives rise to ATG16L1-positive early autophagosome precursors at the plasma membrane, with CLTC knockdown reducing ATG16L1- and LC3-positive puncta (37), while clathrin acting together with AP-1 also supplies a trans-Golgi network-derived membrane source for starvation-induced autophagosome formation (39). The predominantly perinuclear, rather than cortical, distribution of CLTC-LC3/ATG16L1 puncta in our cells is more consistent with this juxtanuclear, Golgi-proximal pool than with a plasma membrane origin. However, our interactome data suggest CLTC is not acting in isolation from the plasma membrane route: LC3 also associated in cytosolic fractions with CLINT1 and the AP-2 subunits AP2A1 and AP2B1, adaptors linked to endosomal and plasma membrane clathrin trafficking, respectively. Together with the ATG16L1 colocalization, this suggests that LC3 engages multiple clathrin-associated trafficking hubs rather than a single membrane source, and that CLTC may serve as a common node coordinating these routes during autophagosome biogenesis. Adding a further layer to this picture, our companion study found that AKT-PH-GFP-positive, phosphoinositide-enriched vesicles partially overlapped with CLTC during cyclic mechanical stretch (45). Together, these observations raise the possibility that CLTC functions as a shared trafficking node linking phosphoinositide-regulated vesicles, autophagosome biogenesis, and nuclear LC3 delivery under mechanical load.

Mechanistically, CLTC may facilitate nuclear transport and LC3 accumulation through several non-mutually exclusive pathways. First, CLTC could promote LC3 nuclear import or retention, potentially through interactions with nuclear transport machinery or scaffolding within nuclear subdomains such as PML bodies. Second, CLTC may stabilize LC3-positive nuclear structures, thereby enhancing their detectability upon inhibition of nuclear export. Third, given its established role in membrane dynamics, CLTC might contribute to the formation of membrane-associated structures within the nucleus or at the nuclear periphery that serve as platforms for LC3 recruitment. A structural, rather than purely vesicular, contribution is plausible: clathrin heavy chain shares a common ancestral coatomer origin with several scaffold nucleoporins, and pharmacological inhibition of clathrin coat assembly acutely disrupts the permeability barrier of the nuclear pore complex itself (46). CLTC may therefore act at the nuclear envelope to modulate LC3 passage as much as within the nucleoplasm to organize LC3-containing structures once inside. This would be consistent with our previous report that nuclear LC3 accumulating in PML bodies under LepB blockade or mechanical stretch (15).

These findings also bear on the physiological role of nuclear LC3 dynamics in TM cells. The TM is a mechanosensitive tissue that continuously adjusts its structure and function in response to fluctuations in intraocular pressure (IOP), and activation of autophagy is now recognized as a central component of this homeostatic response: primary cilia, reciprocal AKT–SMAD2/3 signaling, and, more recently, Class IA PI3Ks and their downstream inositol phosphatases have each been implicated in stretch-induced autophagy in TM cells, and disruption of these pathways impairs the compensatory response to elevated IOP (29, 45). The CLTC-dependent accumulation of nuclear LC3 described here suggests that nuclear LC3 trafficking is itself part of the TM’s adaptive response to mechanical load, rather than an incidental consequence of increased total LC3 expression. Because inappropriate TM adaptation to mechanical stress is thought to contribute to the elevated outflow resistance characteristic of primary open-angle glaucoma, we directly tested whether this nuclear LC3 trafficking arm is altered in gTM cells. nTM and gTM strains showed comparable LepB-induced nuclear LC3-II accumulation under non-stressed conditions, indicating that the basal capacity for nuclear LC3 import is preserved in glaucomatous cells. Mechanical stress, however, substantially blunted this response in glaucomatous cells relative to normal cells, with no changes in CLTC levels observed. This selective, stress-dependent rather than baseline defect suggests that glaucomatous TM cells retain the core molecular machinery for nuclear LC3 trafficking but fail to couple it appropriately to mechanical stress signaling, consistent with a defect in mechanotransduction rather than in autophagy or nuclear transport per se.

Collectively, this work provides the first systematic characterization of the nuclear LC3 interactome in a mechanosensitive primary human cell type and supports a model in which LC3 operates within the nucleus as an organized, compartment-specific interaction network rather than a static reservoir. Within this network, CLTC emerges as a novel regulator of nuclear LC3 accumulation, acting at the interface of vesicle trafficking, nuclear transport, and mechanical stress response. These findings extend the functional repertoire of both LC3 and clathrin beyond their canonical cytoplasmic roles and establish a foundation for future studies of nuclear autophagy-related processes, including nucleophagy and ribophagy, in the context of cellular mechanotransduction and glaucoma pathogenesis.

## CONFLICT OF INTEREST

The authors have declared that no conflict of interest exists.

## ABBREVIATIONS

ACTB: actin beta
AP2A1: adaptor related protein complex 2 subunit alpha 1
AP2B1: adaptor related protein complex 2 subunit beta 1
ATG3: autophagy related 3
ATG4: autophagy related 4
ATG7: autophagy related 7
ATG16L1: autophagy related 16 like 1
BafA1: bafilomycin A1
BSA: bovine serum albumin
CKAP4: cytoskeleton associated protein 4
CLINT1: clathrin interactor 1
CLTC: clathrin heavy chain
CMS: cyclic mechanical stress
CPT1A: carnitine palmitoyltransferase 1A
CYT: cytosolic (fraction)
DAPI: 4’,6-diamidino-2-phenylindole
DMEM: Dulbecco’s modified Eagle medium
DMSO: dimethyl sulfoxide
EPB41: erythrocyte membrane protein band 4.1
EPB41L2: erythrocyte membrane protein band 4.1 like 2
FBL: fibrillarin
FBS: fetal bovine serum
FDR: false discovery rate
FOXK1: forkhead box K1
GABARAP: GABA type A receptor associated protein
GFP: green fluorescent protein
gTM: glaucomatous trabecular meshwork
H2B: histone H2B
HEPES: 4-(2-hydroxyethyl)-1-piperazineethanesulfonic acid
HNRNPK: heterogeneous nuclear ribonucleoprotein K
HSPA8: heat shock protein family A (Hsp70) member 8
IOP: intraocular pressure
IP: immunoprecipitation
LAMP1: lysosomal associated membrane protein 1
LAMP2: lysosomal associated membrane protein 2
LC3/MAP1LC3B: microtubule associated protein 1 light chain 3 beta
LepB: leptomycin B
LIR: LC3-interacting region
MAP1A: microtubule associated protein 1A
MAP1B: microtubule associated protein 1B
MAP1S: microtubule associated protein 1S
Mb Org: membrane/organelle (fraction)
MYH10: myosin heavy chain 10
MYO1C: myosin IC
MYOC: myocilin
NBR1: NBR1 autophagy cargo receptor
NDP52/CALCOCO2: calcium binding and coiled-coil domain 2
NPM1: nucleophosmin 1
NS: non-stressed
nTM: normal (non-glaucomatous) trabecular meshwork
NUC: nuclear (fraction)
OPTN: optineurin
PBS: phosphate-buffered saline
PBS-T: phosphate-buffered saline with Tween 20
PFA: paraformaldehyde
PI3K: phosphoinositide 3-kinase
PLEC: plectin
PML: promyelocytic leukemia (nuclear body)
PVDF: polyvinylidene fluoride
RPL7: ribosomal protein L7
RPL23A: ribosomal protein L23a
SAFB: scaffold attachment factor B
SD: standard deviation
SDS-PAGE: sodium dodecyl sulfate-polyacrylamide gel electrophoresis
siCLTC: CLTC-targeting small interfering RNA
siCNT/siNC: non-targeting control small interfering RNA
siRNA: small interfering RNA
SIRT1: sirtuin 1
SLC25A1: solute carrier family 25 member 1
SQSTM1: sequestosome 1
TM: trabecular meshwork
TP53INP2/DOR: tumor protein p53 inducible nuclear protein 2
TUBA4A: tubulin alpha 4a
WB: western blot
xLIR: expanded/functional LC3-interacting region.

## ACKNOWLEDGEMENTS

The authors acknowledge support from the National Eye Institute (EY026885, EY033600, and EY005722), the BrightFocus Foundation, and Research to Prevent Blindness. The authors also thank Dr. Kate Keller, PhD, of Oregon Health C Science University for providing trabecular meshwork (TM) cells from donors with glaucoma.

## SUPPLEMENTAL FIGURES

**Figure S1. Assessment of purity of the nuclear fractions and co-immunoprecipitation quality**. **(A)** Subcellular fractionation of AdGFP- and AdGFP-LC3–transduced hTM cells. Cytosolic, membrane, and nuclear fractions were analyzed by WB using antibodies against LAMP1, TUBA4A, and Lamin A/C to assess fraction purity. **(B)** Validation of nuclear GFP-LC3 immunoprecipitation. Nuclear extracts were subjected to GFP IP, followed by WB for GFP, SQSTM1 and FBL. Input and IP fractions are shown.

**Figure S2. Proteins differentially associated with nuclear GFP-LC3 in hTM cells following cyclic mechanical stress.** Venn diagram comparing proteins enriched in nuclear GFP-LC3 pulldowns from two independent human trabecular meshwork cell strains, hTM1 and hTM2, under cyclic mechanical stress (CMS) relative to non-stressed conditions (NS), with proteins enriched in AdGFP-LC3 samples relative to AdGFP controls. Proteins in the hTM1 and hTM2 datasets were selected using a CMS-to-NS fold-change threshold of >1.5 and *q* < 0.05. AdGFP-LC3-associated proteins were selected using a fold-change threshold of >1.5, *q* < 0.05, and identification by more than one peptide. Numbers indicate proteins unique to or shared among the three datasets. Proteins within selected overlapping subsets are listed adjacent to the diagram.

**Figure S3. Subcellular distribution of CLTC in nTM and gTM.** Representative immunoblot analysis of cytosolic (CYT), nuclear (NUC), and membrane/organelle (Mb Org) fractions isolated from nTM and gTM cells under non-stressed (NS) and cyclic mechanical stress (CMS) conditions. CLTC was detected in all three subcellular fractions. H2B, LAMP1, and TUBA4A were used as markers for the nuclear, lysosomal/membrane-organelle, and cytosolic fractions, respectively, confirming fraction enrichment. Right, densitometric quantification of CLTC abundance in each fraction, expressed as fold change relative to the corresponding nTM NS condition. No significant differences in CLTC abundance were detected between nTM and gTM cells under either NS or CMS conditions in any fraction. Bars represent mean ± SD, n = 3 independent cell strains experiments.

**Video S1. 3D reconstruction of CLTC and LC3 nuclear interaction**

## Notes

### Competing Interest Statement

The authors have declared no competing interest.

