## Supplemental Figures for "Nuclear LC3 Interactome Profiling Identifies Clathrin Heavy Chain as a Mediator of Nuclear LC3 Translocation in Trabecular Meshwork Cells"

**Figure S1. Assessment of purity of the nuclear fractions and co-immunoprecipitation quality**. **(A)** Subcellular fractionation of AdGFP- and AdGFP-LC3–transduced hTM cells. Cytosolic, membrane, and nuclear fractions were analyzed by WB using antibodies against LAMP1, TUBA4A, and Lamin A/C to assess fraction purity. **(B)** Validation of nuclear GFP-LC3 immunoprecipitation. Nuclear extracts were subjected to GFP IP, followed by WB for GFP, SQSTM1 and FBL. Input and IP fractions are shown.

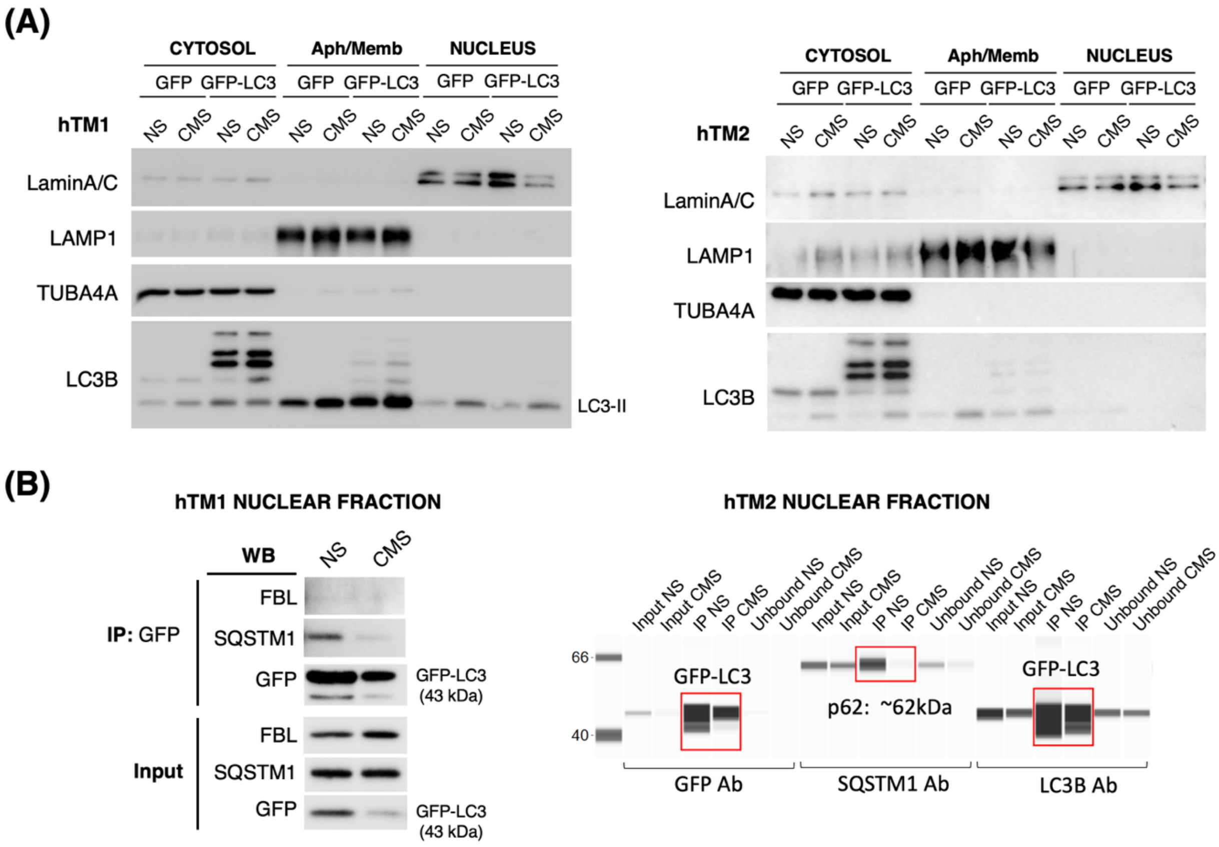

**Figure S2. Proteins differentially associated with nuclear GFP-LC3 in hTM cells following cyclic mechanical stress.** Venn diagram comparing proteins enriched in nuclear GFP-LC3 pulldowns from two independent human trabecular meshwork cell strains, hTM1 and hTM2, under cyclic mechanical stress (CMS) relative to non-stressed conditions (NS), with proteins enriched in AdGFP-LC3 samples relative to AdGFP controls. Proteins in the hTM1 and hTM2 datasets were selected using a CMS-to-NS fold-change threshold of >1.5 and $q < 0.05$. AdGFP-LC3-associated proteins were selected using a fold-change threshold of >1.5, $q < 0.05$, and identification by more than one peptide. Numbers indicate proteins unique to or shared among the three datasets. Proteins within selected overlapping subsets are listed adjacent to the diagram.

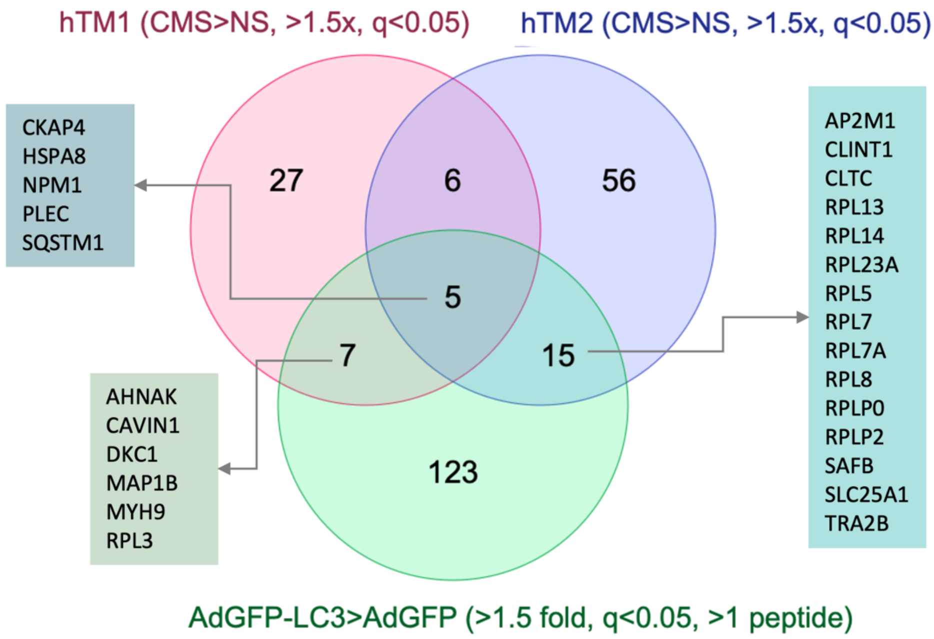

**Figure S3. Subcellular distribution of CLTC in nTM and gTM.** Representative immunoblot analysis of cytosolic (CYT), nuclear (NUC), and membrane/organelle (Mb Org) fractions isolated from nTM and gTM cells under non-stressed (NS) and cyclic mechanical stress (CMS) conditions. CLTC was detected in all three subcellular fractions. H2B, LAMP1, and TUBA4A were used as markers for the nuclear, lysosomal/membrane-organelle, and cytosolic fractions, respectively, confirming fraction enrichment. Right, densitometric quantification of CLTC abundance in each fraction, expressed as fold change relative to the corresponding nTM NS condition. No significant differences in CLTC abundance were detected between nTM and gTM cells under either NS or CMS conditions in any fraction. Bars represent mean ± SD, n = 3 independent cell strains experiments.

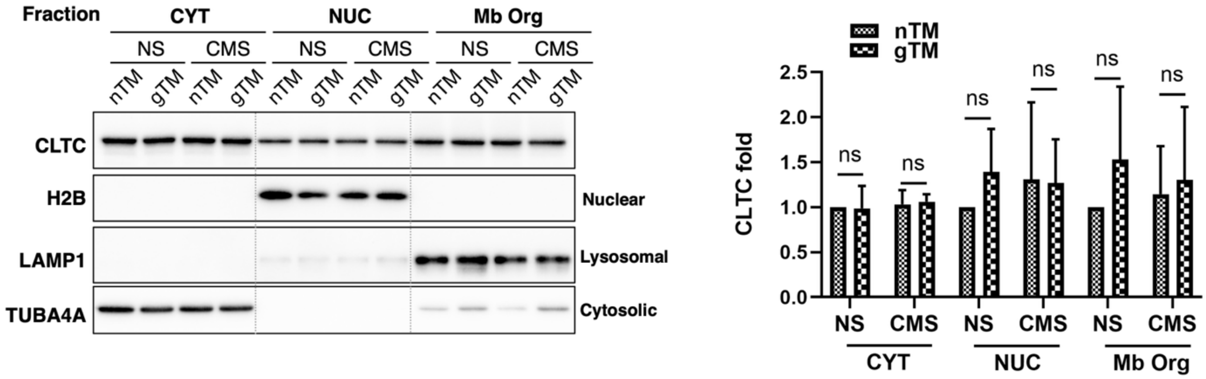
