## Supplementary figures and images for "Nuclear LC3 Interactome Profiling Identifies Clathrin Heavy Chain as a Mediator of Nuclear LC3 Translocation in Trabecular Meshwork Cells"

### Supplemental Videos

#### Slide 1
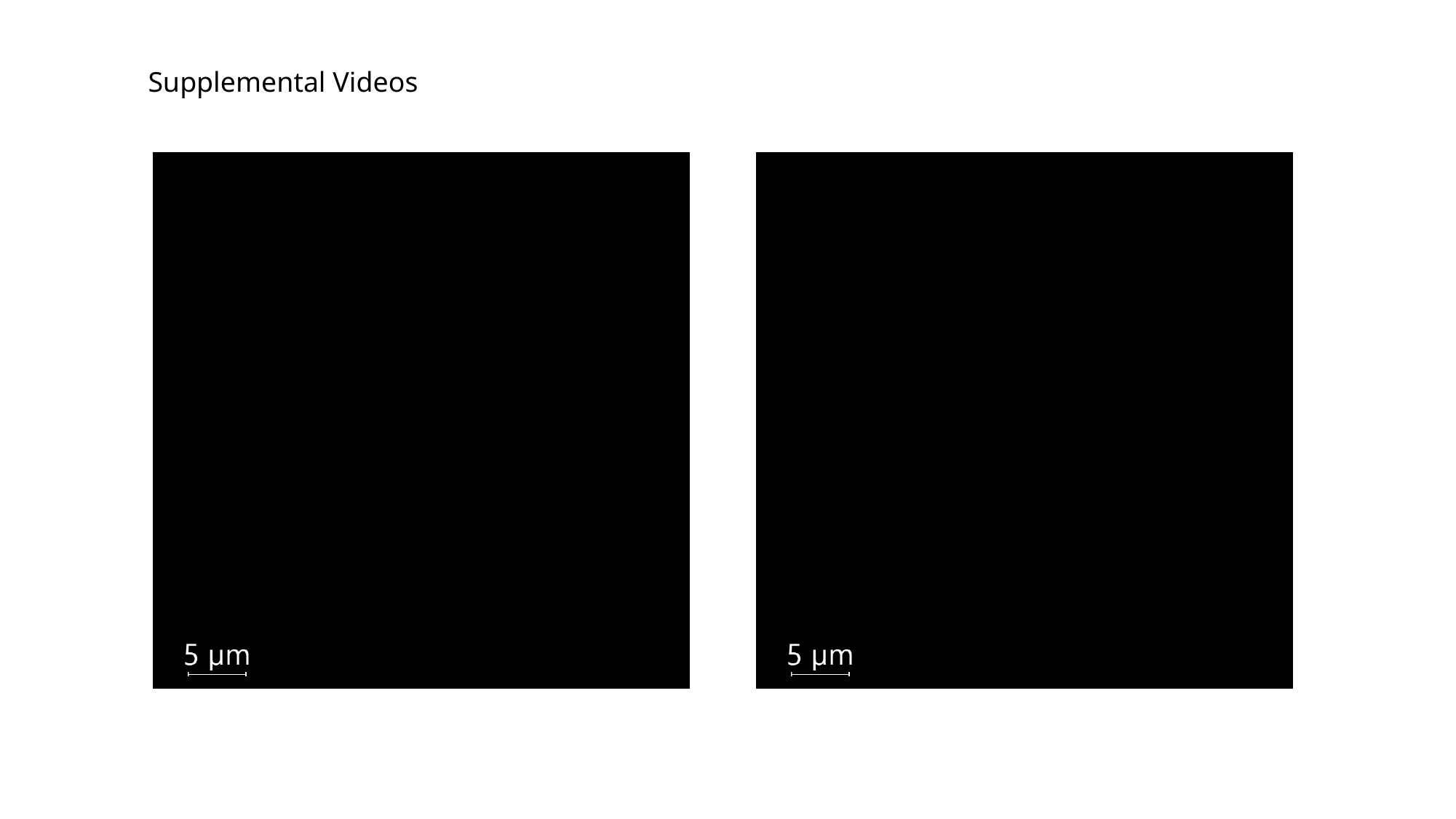

Supplemental Videos
